# Novel mouse model reveals neurodevelopmental origin of PMM2-CDG brain pathology

**DOI:** 10.1101/2025.06.01.657261

**Authors:** Andrew C. Edmondson, Rohit Budhraja, Zijie Xia, Ashley Melendez-Perez, Cadmus Cai, Silvia Radenkovic, Ashley M. Collins, Emily J. Shiplett, Sophie F. Hill, Ala Somarowthu, Johanna Dam, Ling-Lin Pai, Mariarita Santi, Seonhee Kim, Miao He, Ethan M. Goldberg, Tamas Kozicz, Eva Morava, Akhilesh Pandey, Zhaolan Zhou

## Abstract

Congenital disorders of glycosylation (CDG) are a group of neurogenetic conditions resulting from disruptions in the cellular glycosylation machinery. The majority of CDG patients have compound heterozygous pathogenic variants in the phosphomannomutase 2 (*PMM2)* gene. Individuals with PMM2-CDG exhibit multi-systemic symptoms, prominently featuring neurological deficits with nearly all patients exhibiting cerebellar hypoplasia and ataxia. To overcome embryonic lethality caused by whole body knock-out of *Pmm2* and mimic patient-related compound heterozygous pathogenic variants, we paired a *Pmm2* flox allele (*Pmm2*^fl^) with a catalytically inactive knock-in allele (*Pmm2*^R137H^), commonly present in PMM2-CDG patients. Mice with post-mitotic loss of PMM2 from neurons or astrocytes are indistinguishable from unaffected littermates, including in a broad battery of neurological assessments. In contrast, removal of PMM2 from embryonic neural precursor cells leads to cerebellar hypoplasia, ataxia, seizures, and early lethality. Comprehensive multi-omics profiling, including metabolomics, glycomics, single-cell transcriptomics, proteomics, and glycoproteomics, reveal widespread molecular disturbances throughout the brain, with the cerebellum showing the most pronounced disruption. These findings highlight the heightened dependency of the developing cerebellum on intact N-glycosylation, aligning with clinical observations in PMM2-CDG patients. Importantly, glycoproteomic alterations identified in our mouse model are corroborated in PMM2-CDG patient post-mortem cerebellar tissue, underscoring the translational relevance of our findings and implicating impaired synaptic transmission as a key pathogenic mechanism.

## Introduction

Congenital disorders of glycosylation (CDG) are a group of multi-system genetic disorders that disrupt cellular glycosylation machinery. The most common CDG, PMM2-CDG, is caused by biallelic pathogenic variants in *PMM2*, a gene encoding the phosphomannomutase 2 (PMM2) enzyme necessary for N-linked protein glycosylation^1–3^. PMM2-CDG has an estimated incidence of approximately 1:35,000 in North America and Europe, in part due to high carrier status of a common pathogenic variant, R141H. Patients with PMM2-CDG typically suffer from multi-systemic symptoms, including growth failure, coagulopathies, and endocrinopathies^2–4^. Even after stabilization of symptoms in other organs, PMM2-CDG patients exhibit cerebellar hypoplasia (particularly of the cerebellar vermis)^5^, ataxia^2–7^, and other neurological deficits, including epilepsy^2–4^. Pathophysiology of PMM2-CDG is thought to originate from protein hypoglycosylation^8^, but the connection between disruption of protein glycosylation and patient symptoms (especially of glycoproteins responsible for neurological symptoms) has largely not been determined. Recent reports in human brain organoid^9^ and zebrafish^10^ models of PMM2-CDG have also implicated pathogenic rewiring of metabolic pathways as potential contributors to disease symptoms^11,12^.

The genetic basis of PMM2-CDG provides an opportunity to investigate its pathophysiology using mouse models; however, current models fail to recapitulate the neurological features of PMM2-CDG^13–15^. Four mouse alleles have been reported in prior attempts to model PMM2-CDG, including a knockout allele that exhibited early embryonic lethality in homozygosity^14^ and three knock-in alleles of single amino acid changes, including alleles equivalent to the two most prevalent human pathogenic variants R141H (NM_000303.2:c.422G>A (p.Arg141His), ClinVar 7706) and F119L (NM_000303.2:c.357C>A (p.Phe119Leu), ClinVar 7711)^13,15^. The hypomorphic alleles generated to date have either so mildly reduced PMM2 enzymatic activity that the mice have not developed a discernable phenotype^15^, or have so severely reduced PMM2 enzymatic activity that they result in embryonic lethality^13–15^. While this highlights the importance of glycosylation, particularly during embryonic development, it presents a challenge to modeling the disorder in mice. It is also possible that some organ-specific reliance on N-glycosylation in the developing mouse differs from humans, such that the degree of reduction of PMM2 enzyme activity to produce neurological pathology exceeds a threshold that is compatible with life in other organs. Consistent with this, a recently reported compound heterozygous mouse model combining the two most common human alleles continued to exhibit some embryonic lethality (with fewer than expected compound heterozygous pups born) and surviving pups exhibited severe systemic symptoms resulting in early lethality^13^. Even the few surviving mice they were able to evaluate failed to recapitulate the neurological pathology of PMM2-CDG^13^.

Given these challenges, we leveraged a conditional *Pmm2* mouse flox allele (*Pmm2*^fl^) developed as part of the European conditional mouse mutagenesis (EUCOMM) program^16^ to enable cell-type specific severe reduction of PMM2 enzymatic activity while avoiding embryonic lethality. We paired *Pmm2*^fl^ with the previously developed catalytically inactive *Pmm2*^R137H^ allele^13^ (equivalent to R141H, a human pathogenic variant carried by more than 60% of PMM2-CDG patients^2^) to recapitulate the compound heterozygous state of PMM2-CDG individuals and incorporate spatial-temporal control over PMM2 activity dependent on the presence of Cre recombinase. Given the prominent neurological concerns of PMM2-CDG patients, we focused on PMM2 in the brain and generated neural knock-out models of *Pmm2* using a series of brain-specific Cre lines. In an effort to dissect cell-type contributions to PMM2-CDG neurological pathology, we initially used pan-neuronal post-mitotic Cre (Snap25-IRES2-Cre-D^17,18^, nKO) and astrocyte-specific Cre (Tg(Gfap-Cre)^19^, aKO) lines, but subsequently also utilized Nestin-Cre^20^ (eKO), which expresses Cre during mid-embryonic development in neuronal and glial cell precursors. These efforts produced mice with drastically different phenotypes, revealing a neurodevelopmental pathogenic mechanism to PMM2-CDG brain pathology and resulting in the first PMM2-CDG mouse model with prominent cerebellar manifestations.

## Results

### Molecular validation

We utilized a conditional flox allele of *Pmm2* (*Pmm2*^fl^) generated by the European Conditional Mouse Mutagenesis (EUCOMM) Program^16^, in which exon 3 of *Pmm2* is flanked by *LoxP* sites and upon Cre-mediated excision of exon 3 a reading frameshift leads to a premature stop codon, p.V48Efs*24, resulting in the loss-of-function allele *Pmm2*^KO^ (**Fig. 1A**). The loss of function nature of this allele was validated by the International Mouse Phenotyping Consortium, where homozygosity for the *Pmm2*^KO^ allele results in embryonic lethality^21^. We also obtained a carrier of the *Pmm2*^KO^ allele as the result of leaky germline Cre expression^22^, which was backcrossed on the C57Bl/6 background and the resulting carrier offspring were bred together. We found that 41 pups genotyped at postnatal (P) day 0 and 1 from 7 different litters failed to produce any homozygous KO mice with 36.6% (n = 15) of pups being wild-type and 63.4% (n = 26) of pups being Cre-recombined allele carriers (*P* = 0.0009, χ^2^ test), consistent with previous studies of loss-of-function alleles of *Pmm2*^13–15^.

**Fig 1.**
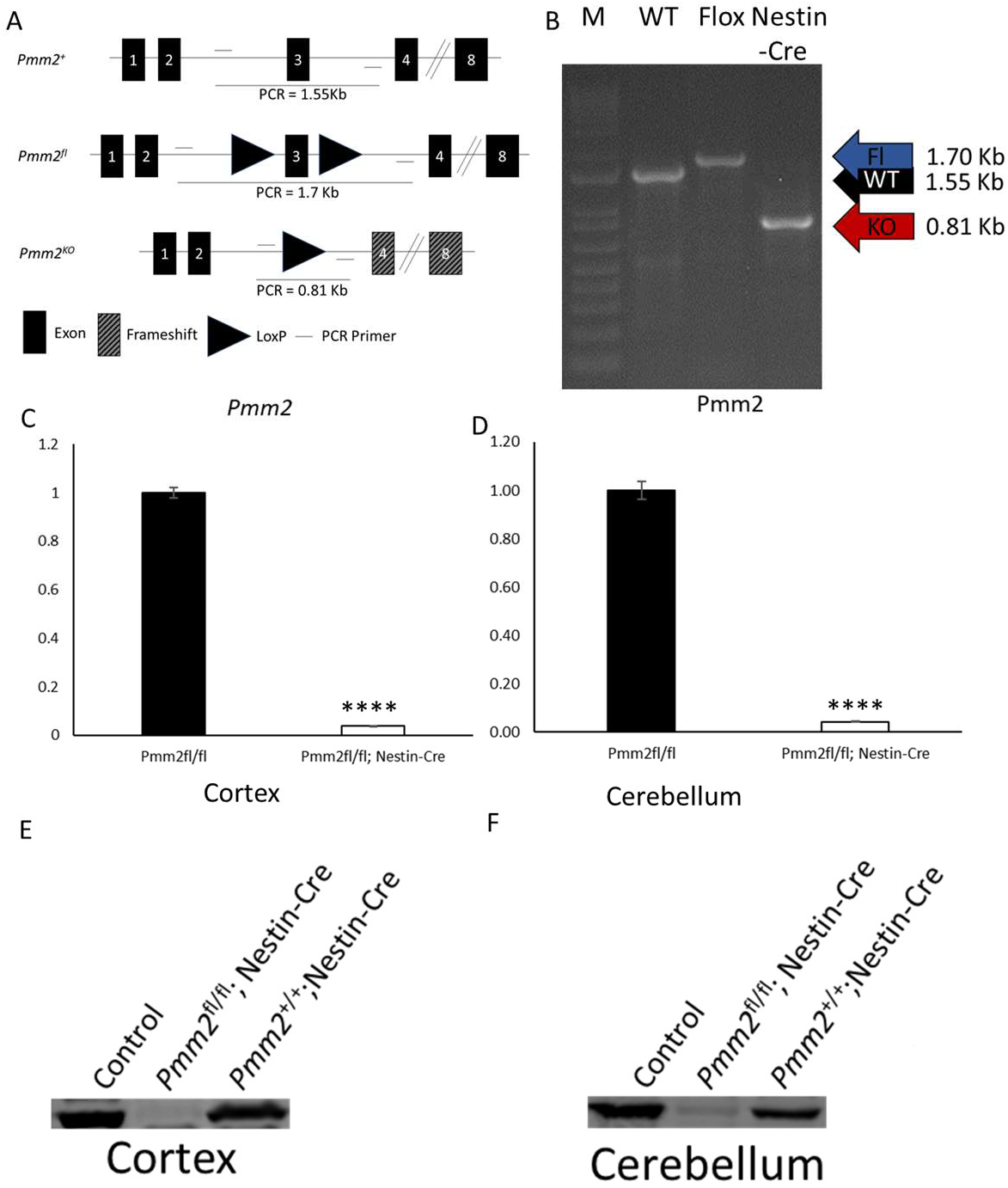
Molecular validation of *Pmm2* eKO mice. A, Schematic of mouse genomic loci of *Pmm2^fl^* (conditional KO) and *Pmm2^KO^* (KO) alleles with genomic recombination PCR assay strategy indicated. B, Genomic Recombination Assay PCR results of cortical DNA from WT, Conditional KO (Flox) and eKO (Cre) mice. C and D, qPCR quantitation of *Pmm2* expression in cortex and cerebellum, respectfully. E and F, Western for PMM2 protein in cortex and cerebellum, respectfully. Welch’s t-test. ****, P < 0.0001

For the majority of studies, to prevent germline recombination of the *Pmm2*^fl^ allele from ectopic germline Cre expression, we crossed *Pmm2*^R137H/+^ mice with the different Cre lines and bred resulting mice carrying both *Pmm2*^R137H/+^ and Cre to *Pmm2*^fl/fl^ mice in order to generate compound heterozygote mice where one allele of *Pmm2* is floxed and one allele bears the R137H point mutation (*Pmm2*^R137H/fl^) for experimental studies. Snap25-Cre and Tg(Gfap-Cre) alleles were maintained in the female mice due to leaky germline expression of Cre in male mice^22^. However, for molecular validation studies in cortex and cerebellum, given that the Pmm2^R137H/+^ allele generates a stable transcript and catalytically inactive protein, we initially generated homozygous *Pmm2*^fl/fl^ mice with and without Cre. Molecular validation studies of cortex and cerebellum included a polymerase chain reaction (PCR) based genomic recombination assay to confirm Cre-mediated recombination of LoxP sites and excision of exon 3, quantitative PCR (qPCR) to quantify *Pmm2* expression, and Western blot to evaluate PMM2 expression. Our eKO mice demonstrated robust genomic recombination (**Fig. 1B**), decreased *Pmm2* expression (**Fig. 1 C and D**) and decreased PMM2 protein (**Fig. 1 E and F**). Molecular validation studies in nKO mice (**Supp. Fig. 1**) and aKO mice (**Supp. Fig. 2**) were consistent with robust cell-type specific Cre-mediated recombination occurring as desired in only post-mitotic neurons or astrocytes respectively, with the astrocyte-specific PMM2 contribution in brain being significantly less than from neurons.

### Biochemical Characterization

The PMM activity of PMM2 is responsible for reversibly converting mannose-6-phosphate (Man-6-P) to mannose-1-phosphate (Man-1-P), the precursor to GDP-Mannose (GDP-Man), an essential building block of glycosylation. The efficient Cre-mediated recombination observed in our mouse models (**Fig. 1B** and **Supp. Figs. 1A** and **2A**) would be expected to result in cells with both a catalytically inactive (R137H) *Pmm2* allele and a frameshift loss-of-function *Pmm2* allele and concomitant drastic reduction in cytosolic PMM2 enzyme activity. We utilized an established phosphomannomutase (PMM) enzyme activity assay used to clinically diagnose PMM2-CDG patients (**Fig. 2A**)^23^ to biochemically measure PMM enzyme activity in cortex and cerebellum of our mouse models. Of note, performed on bulk brain lysate, the measured PMM enzyme activity additionally includes catalytic activity of PMM2 in non-Cre-recombined cells (such as cell populations in the brain excluded by the use of cell-type specific Cre drivers), as well as of other related enzymes (such as phosphomannomutase 1 (PMM1) and phosphoglucomutase 1 (PGM1)) with *in vitro* PMM activity^23,24^. With this in mind, we confirmed that brains from nKO and eKO mice exhibit significantly reduced PMM enzymatic activity (**Fig. 2-D**) and, consistent with molecular validation studies, observed that astrocytes contribute a minor portion of PMM2 activity to bulk brain lysate (**Fig. 2E**). Notably, PMM activity was similar within the cortex and cerebellum of each model with the amount of PMM activity differing across models likely due to age-related abundance differences of PMM2 and other contributing enzymes in the brain (eKO model was assayed at P12, the nKO and aKO models were assayed at ∼P120). Both eKO and nKO demonstrated similar reductions in PMM enzyme activity, displaying approximately 50% PMM enzyme activity of their littermate controls, respectively.

**Fig. 2.**
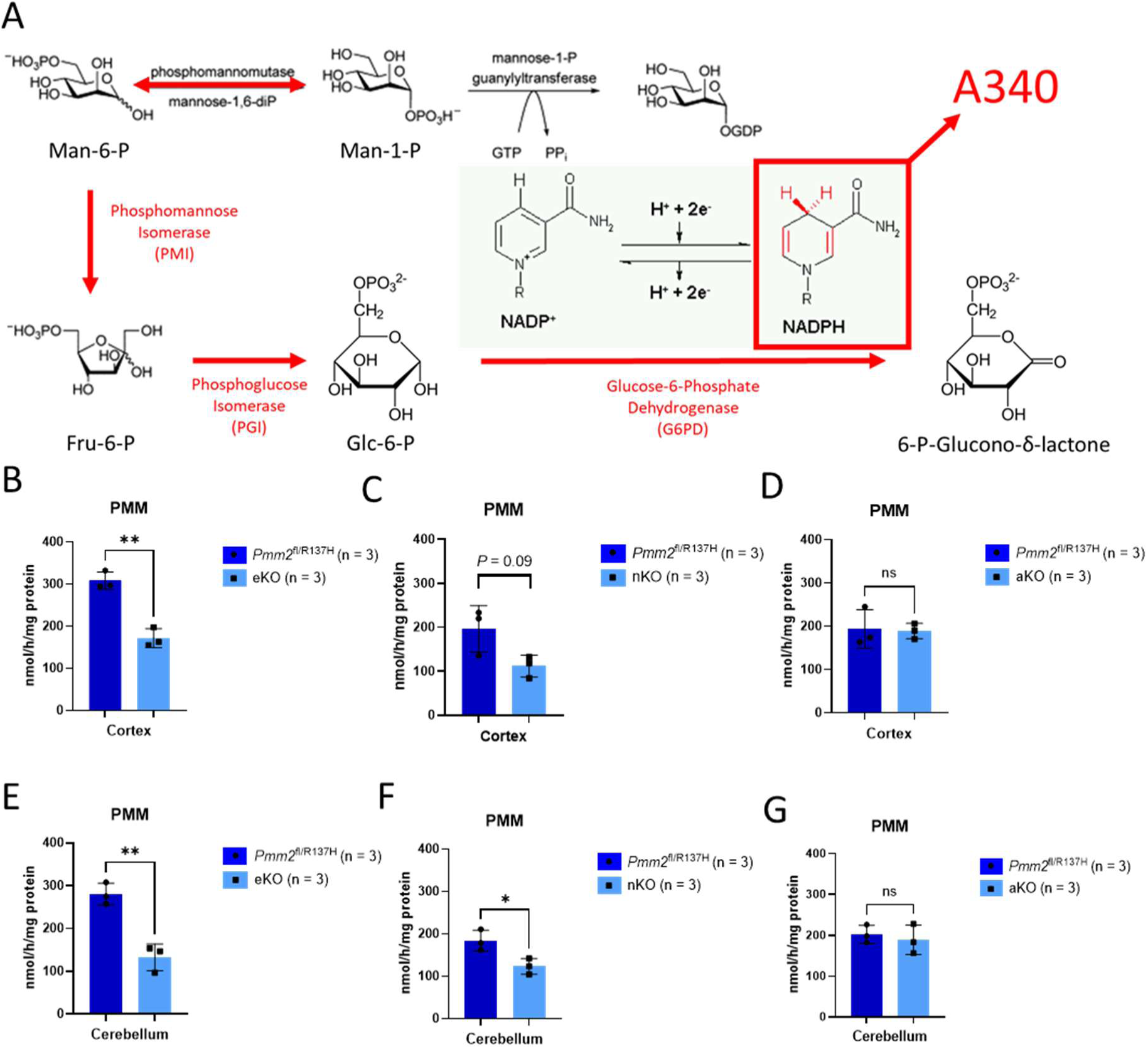
Brain PMM enzyme activity. (A) Overview of biochemical assay used to measure PMM activity where added Man-1-P is metabolized to Glucose-6-Phosphate (Glc-6-P) by exogenous MPI and PGI which is then converted to 6-P-Glucono-delta-lactone by exogenous G6PD through a NADP-dependent reaction and changes in NAPDH abundance are reflected in light absorbance at 340 nm. PMM enzyme activity in cortex in (B) eKO, (C) nKO, (D) aKO and cerebellum in (E) eKO mice, (F) nKO, and (F) aKO. Data are represented as mean ± SD. Welch’s unpaired, 2-tailed t-test. *, *P* < 0.05, **, *P* < 0.01.

### Birth Ratios and Growth

Mice were ear tagged and genotyped via tail biopsy between P12-P14. Genotypes across litters were monitored and demonstrated no evidence of embryonic lethality in affected mice for any of the models. For mice with Nestin-Cre, 57 pups across 9 litters yielded 28% affected eKO mice (*P* = 0.94, χ^2^ test). For mice with Snap25-Cre, 79 pups across 12 litters yielded 26% nKO mice (*P* = 0.7, χ^2^ test). For mice with Tg(Gfap-Cre), 99 pups across 15 litters yielded 20% aKO mice (*P* = 0.6, χ^2^ test).

Mice were weighed weekly starting at weaning and monitored twice weekly to assess for neurological phenotypes. For nKO and aKO mouse models, both male and female mice demonstrated normal weight gain relative to unaffected littermates (**Supp. Fig. 3**). However, it soon became apparent that eKO mice rarely survived to P21 (**Fig. 3B**). Closer evaluation of eKO mice identified that a rapid oscillatory whole-body shaking phenotype emerged around P9 to P10 and that as unaffected littermates subsequently learned to walk, eKO mice failed to perfect their gait and exhibited ataxic ambulation with frequent loss of righting reflex. They are smaller than unaffected littermates (data from P14 after ear tagging shown in **Fig. 3A**) and fail to gain weight. Starting around P16 and with increasing frequency on following days, affected eKO mice demonstrated abnormal movements, including pausing behaviors, tonic stiffening, and hindlimb extension concerning for seizures. While eKO mice typically recovered from these episodes when observed, occasionally the mice were observed to die following these behaviors, although most deaths occurred unobserved.

**Fig. 3.**
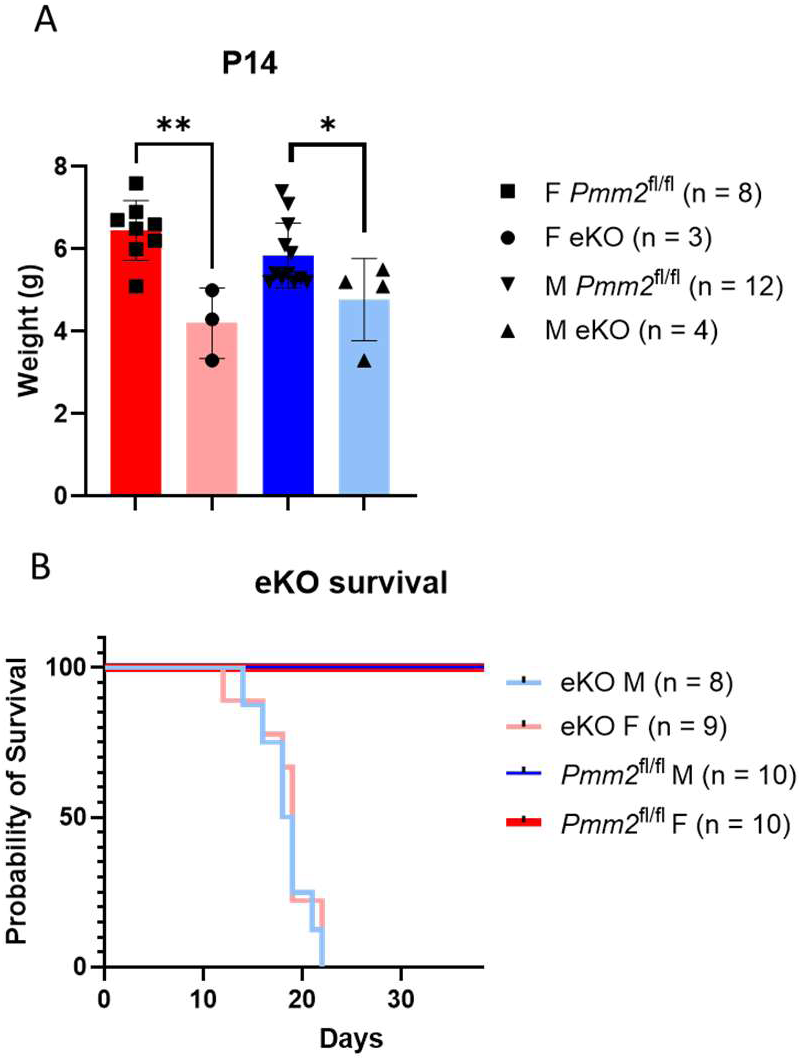
*Pmm2* eKO mice are small and die prematurely. A) Weights of female and male mice at P14, grouped by genotype. C) Kaplan-Meier survival curve of eKO mice. Data are represented as mean ± SD. Welch’s unpaired, 2-tailed t-test. Number of biological replicates indicated in parentheses. *, *P* < 0.05; **, *P* < 0.01.

In an attempt to determine the cause of death, a complete clinical veterinary pathological assessment via necropsy was completed on 6 affected eKO (3 males, 3 females) and 6 unaffected littermates (2 males, 4 females) between P18-P20 days of age. A definitive cause of death could not be identified. Affected mice were noted to have decreased adipose stores but without histologic pathology (data not shown), suggesting possible decreased ability to feed due to ataxia. Affected mice were also noted to have prominent renal tubular cysts occupying approximately 50% of the volume of each renal cortex (data not shown). The renal findings were considered attributable to the use of Nestin-Cre, which has known ectopic expression in the renal tubule cells of the kidneys and remaining tubules as well as other components of the kidney were histologically unremarkable, suggesting an important role of PMM2 and N-glycosylation in renal development. Cystic kidneys have also been reported in some PMM2-CDG patients^25^. Serum electrolytes and creatine were measured and normal (data not shown), suggesting that the affected kidneys appear to retain enough functional perimortem capacity, although their reserve capacity is likely decreased.

Clinical pathology evaluation noted no major pathologic changes in the CNS and the functional neurologic defects were thought to be the most likely cause of death, given the clinical signs, lack of detectable structural/morphological changes in the CNS, and lack of severe lesions elsewhere in the body (data not shown).

### Histologic Evaluation of eKO Brain

Given the abnormal ataxia in eKO mice and correlate with PMM2-CDG patients, we pursued further histologic evaluation of the eKO cerebellum, including Calbindin staining to evaluate Purkinje cells. The cerebellum was significantly smaller in *Pmm2* eKO mice at both P10 (**Supp. Fig. 4A, B**) and P17 (**Fig. 4A, B**), with evidence of vermian hypoplasia with a reduced midsagittal cerebellum area relative to unaffected controls for both timepoints (**Supp. Fig. 4C, D**; **Fig. 4C, D**).

**Fig 4.**
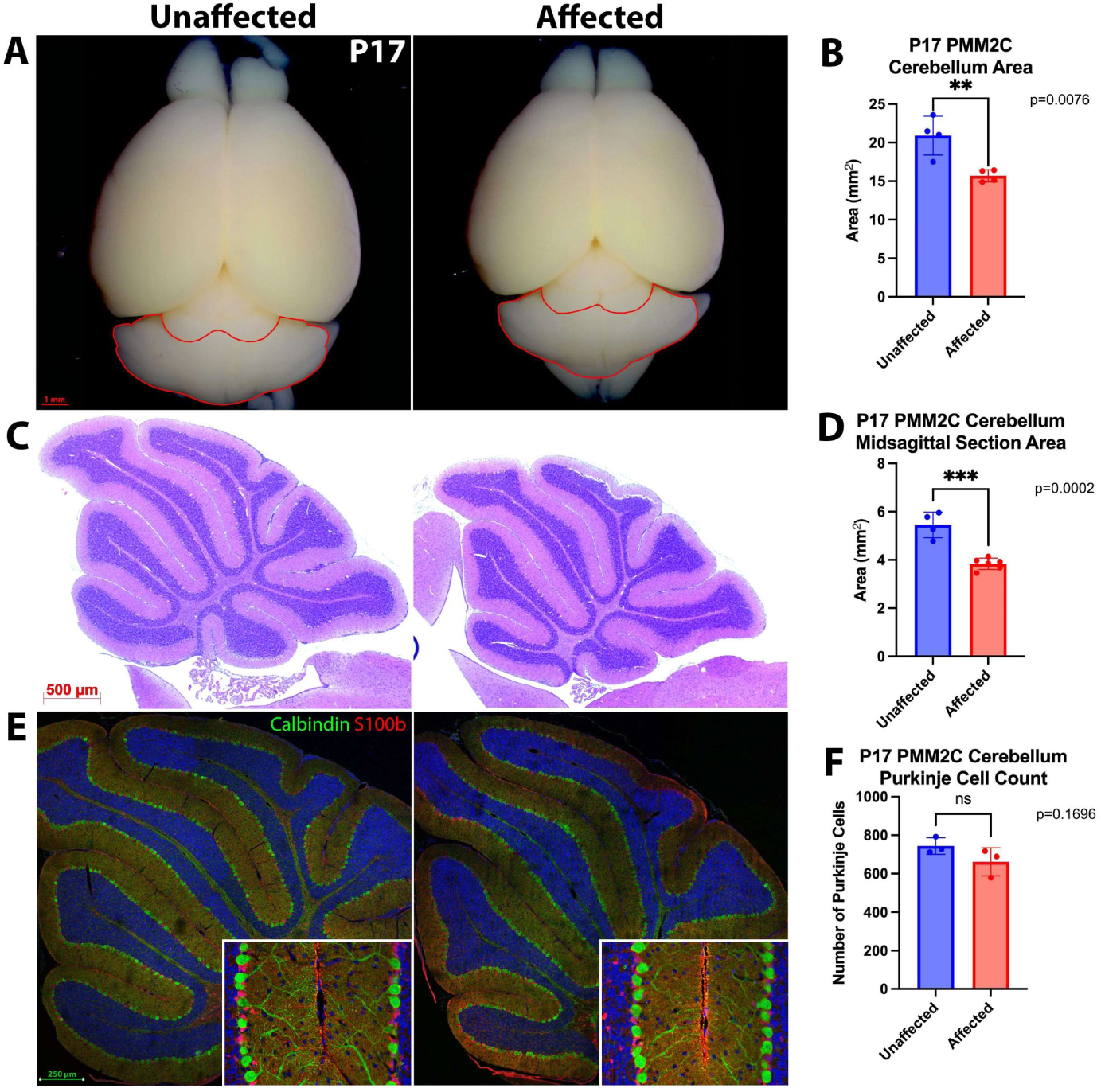
Histologic evaluation of cerebellum in P17 *Pmm2* eKO mice. A, Cerebellar area of unaffected (*Pmm2*^fl/+^; Nestin-Cre) and affected (*Pmm2*^R137H/fl^; Nestin-Cre) mouse brains at P17. B, Unaffected and affected P17 mice show a significant difference in cerebellum area, *P* = 0.0076, n = 4 for each genotype. C, Representative images of midsagittal sections stained with hematoxylin and eosin of P17 unaffected and affected mice. D, Unaffected and affected P17 mice show a significant reduction in midsagittal area, *P* = 0.0002, unaffected n = 4, affected n = 6. E, Representative immunohistochemistry images of cerebellum of unaffected and affected mice with staining for Purkinje cells (Calbindin), glia (S100b), and DNA (Hoechst). F, Purkinje cell numbers in the unaffected and affected P17 mice remain unchanged between the two groups. Comparisons analyzed using an unpaired student’s t-test, plotted as mean ± SD. **, *P* < 0.01; ***, *P* < 0.001.

Notably, despite the hypoplasia, affected brains appeared structurally normal, and quantification of Purkinje cells revealed no significant differences between unaffected and affected mice at neither P10 nor P17 (**Suppl Fig. 4E, F**; **Fig. 4E, F**). We also did not identify any overt structural abnormalities in the brains of affected mice. However, at P10, affected mice exhibited a modest but statistically significant reduction in brain size, as measured by hemispheric area. Despite this decrease, cortical layers appeared intact, with no observable differences in cortical thickness nor the distribution of CTIP2+ (layer V) and Cux1+ (layers II–IV) cells compared to unaffected controls (**Supp. Fig. 5**). These evaluations suggest the abnormal neurologic signs in eKO mice are primarily due to functional rather than structural defects.

### In Vivo EEG Evaluation

To further identify and characterize the abnormal antemortem movements and evaluate for functional neurological deficits, we performed *in vivo* EEG monitoring of affected mice and littermate controls between ages P17-P18 and captured numerous seizures in affected mice. Four of the 5 affected mice recorded demonstrated seizures during the 6-8.5 hour recording window. The affected mice had an estimated average of 1.23 ± 0.49 seizures/hr and average seizure duration of 1.41 ± 0.7 mins over the recording period. Of note, the affected mouse not observed to have seizures was the youngest of the recorded mice, supportive of the observation that the seizures increase in frequency and duration as the mice age (**Fig. 5**). These EEG measurements of brain activity document functional defects at the cellular or circuit level in *Pmm2* eKO mice, supportive of a disruption in neurotransmission underlying pathogenesis in PMM2-CDG.

**Fig. 5.**
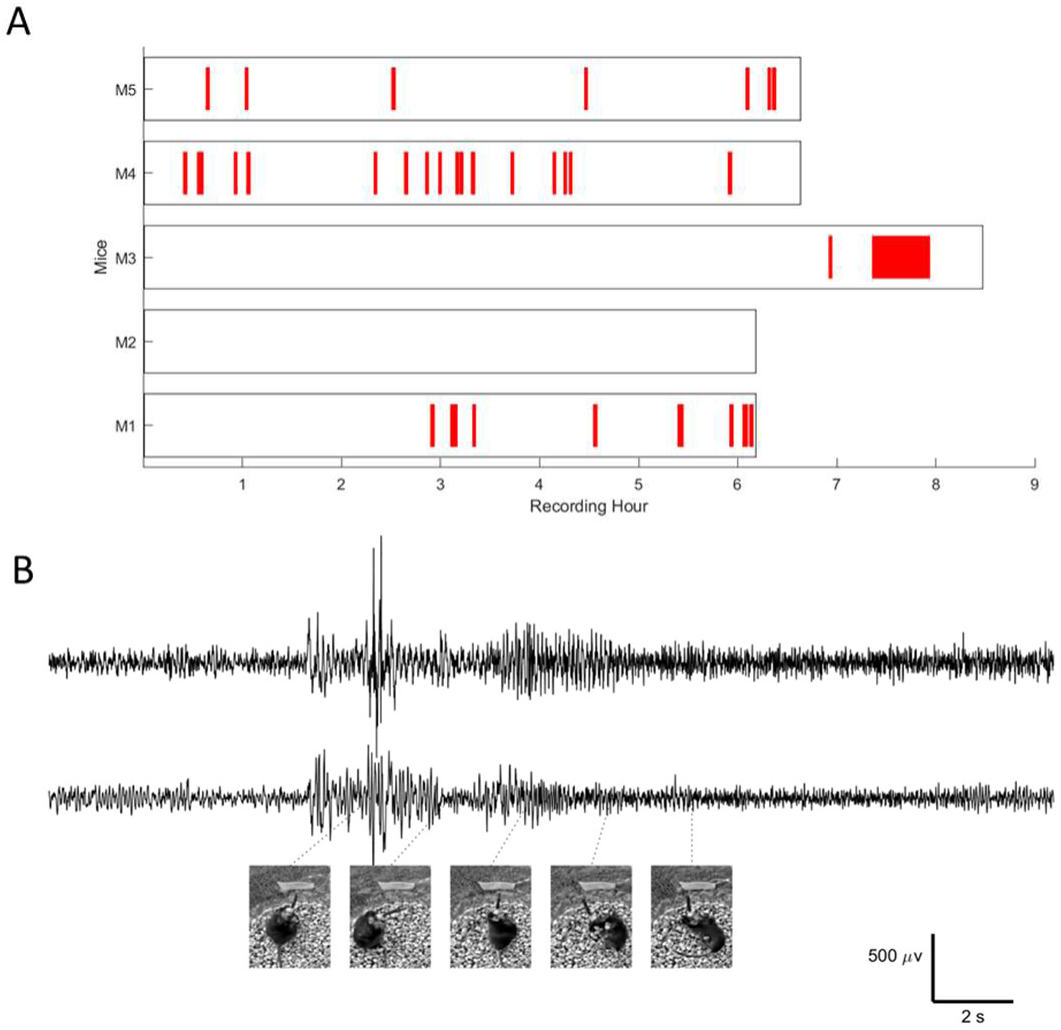
In vivo EEG recording in *Pmm2* eKO mice identify seizures. (A) Raster plot of recordings in affected mice with episodes of seizures indicated (red bars). (B) Representative EEG trace from a typical seizure in mouse M4 at P18 with behavioral poses from the identified timepoints throughout the seizure. μV and time scale bars indicated.

### Behavioral Evaluation

To systematically characterize the contribution of neuronal and astrocyte PMM2 to PMM2-CDG relevant behavioral phenotypes we performed a battery of behavioral tests over a 2-month period starting at 10-12 weeks of age, however nKO and aKO mice failed to demonstrate behavioral deficits compared with littermate controls (data not shown). Clinical histopathologic analysis of the brains from nKO and aKO mice were also normal (data not shown).

### Metabolomics

Given the lack of relevant neurologic finding in nKO and aKO mice, further studies focused on the eKO mice. We employed a comprehensive multi-omic approach to characterize eKO mice and obtain molecular insight into the pathogenic mechanisms underlying their striking phenotypes. In order to identify proximal deficits rather than perimortem secondary changes, omics-based evaluations were performed in P12 mice, when the mice were clearly and reliably exhibiting ataxic behavior, but before they had begun to exhibit behavioral seizures, or earlier. The metabolomic evaluation demonstrated evidence consistent with biochemical blockade at the level of PMM2 with elevations of hexose-phosphate pool (likely due to elevations Man-6-P) (**Fig. 6C**) and decreases of GDP-Man (**Fig. 6D**).

**Fig. 6.**
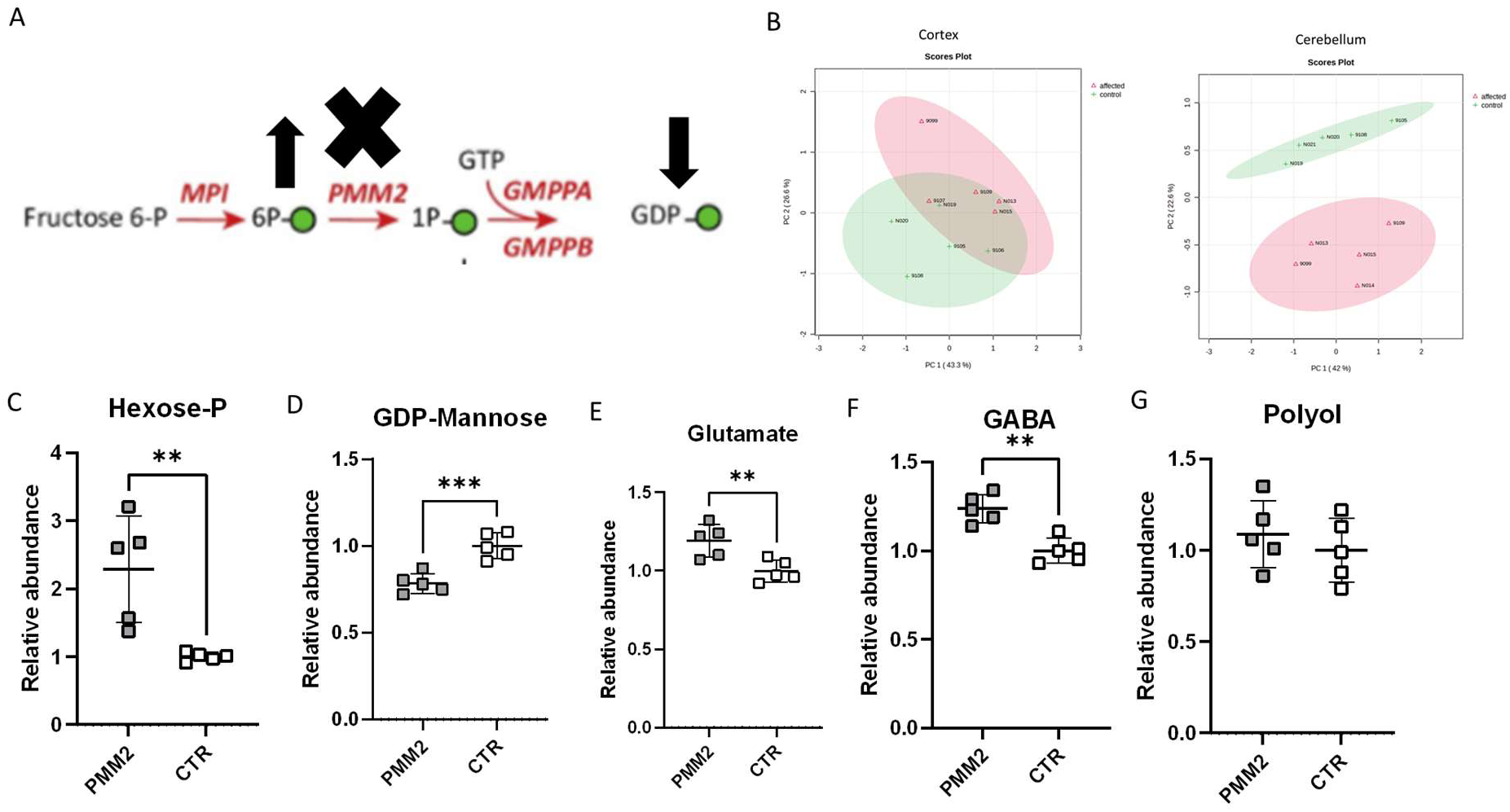
Metabolomic alterations in eKO cerebellum. Overview of PMM2 biochemical pathway illustrated in (A). (B) Principal components analysis of metabolomic data. Select biochemical metabolites of particular interest (C) Hexose-P (likely representing increased Man-6-P), (D) GDP-Man, (E) neurotransmitter glutamate, (F) GABA, (G) polyol pool (including sorbitol among other polyols). Welch’s unpaired, 2-tailed t-test. **, *P* < 0.01, ***, *P* < 0.001. Individual data points are plotted with line representing mean ± SD.

Interestingly, principal components analysis suggested the eKO mice had more pronounced metabolic abnormalities in the cerebellum than in the cortex (**Fig. 6B**). We also saw evidence of elevated neurotrasmitters, glutamate and gamma-aminobutyric acid (GABA) (**Fig. 6 E and F**), highlighting a role for altered synaptic neurotransmission in PMM2-CDG brain pathogenesis. Of note, the polyol pool (which contains sorbitol) was not altered in brain (**Fig. 6G**). Alternatively, rather than shunting accumulating Man-6-P through sugar alcohol reductase pathways, we observed supportive evidence of increased metabolites in hexosamine biosynthesis and the pentose phosphate pathway (**Supp. Fig. 10**).

While less prominent metabolomic alterations were evident in cortex, the majority of the metabolic alterations overlap in cortex and cerebellum with similar trends in direction of change even when not statistically significant in cortex (data not shown).

### Glycomics

We performed N-glycomic evaluation of the cerebellum at P12. Consistent with decreased PMM2 enzymatic activity and the metabolic blockade leading to elevations of Man-6-P and deficiency of GDP-Man, we identified an accumulation of smaller oligomannose N-glycans and a decrease in larger oligomannose N-glycans, consistent with changes typically seen in PMM2-CDG individuals (**Fig. 7**).

**Fig. 7.**
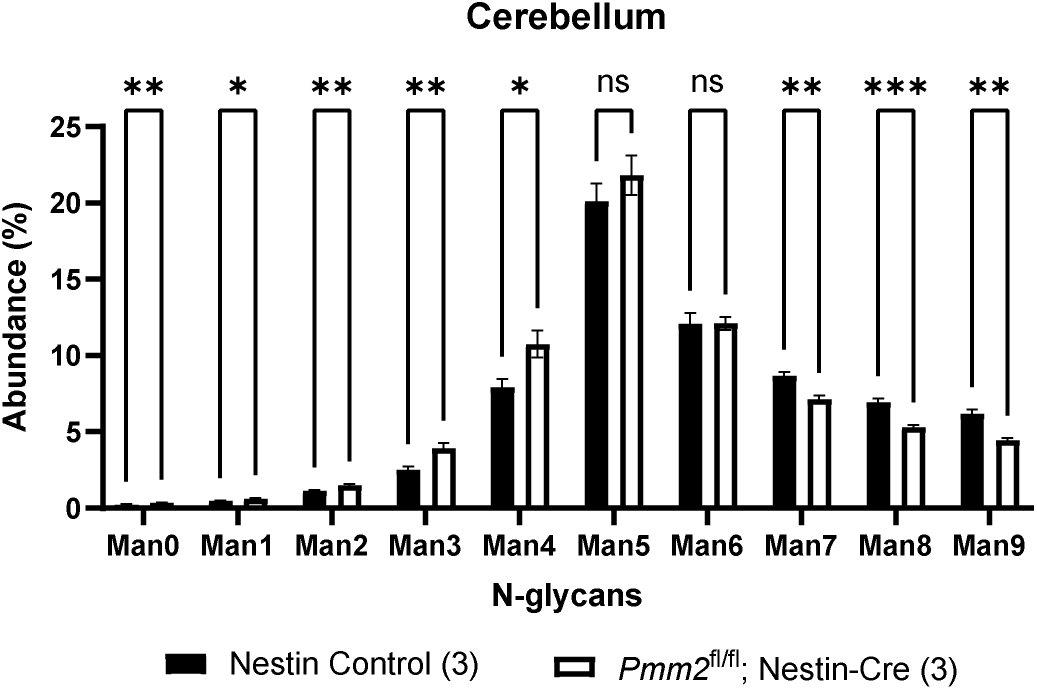
Cerebellum glycomics demonstrate evidence of GDP-mannose deficiency consistent with PMM2-CDG. N-glycan species abundance in cerebellum. Number of samples indicated in paranthesis. Welch’s unpaired, 2-tailed t-test. *, *P* < 0.05, **, *P* < 0.01, ***, *P* < 0.001. Data are represented as mean ± SD.

### Single-cell transcriptomics

In order to identify proximal transcriptomic deficits rather than pathology-driven secondary transcriptional changes, single cell transcriptomics was performed in cortex and cerebellum samples from P7 mice (n = 4 of each genotype), prior to the emergence of shaking or ataxic behavior (**Fig. 8**). We observed dysregulated transcriptional pathways replicated in similar patterns of across multiple cell clusters in both cortex and cerebellum. We observed transcriptional upregulation of genes related to unfolded protein response (UPR) and ER stress pathways, as well as related to oxidative phosphorylation and mitochondrial function, suggestive of mitochondrial dysfunction and impaired glucose metabolism. We also observed transcriptional downregulation of neurodevelopmental genes associated with neurogenesis, neuronal differentiation, axon guidance, and synapse organization.

**Fig. 8.**
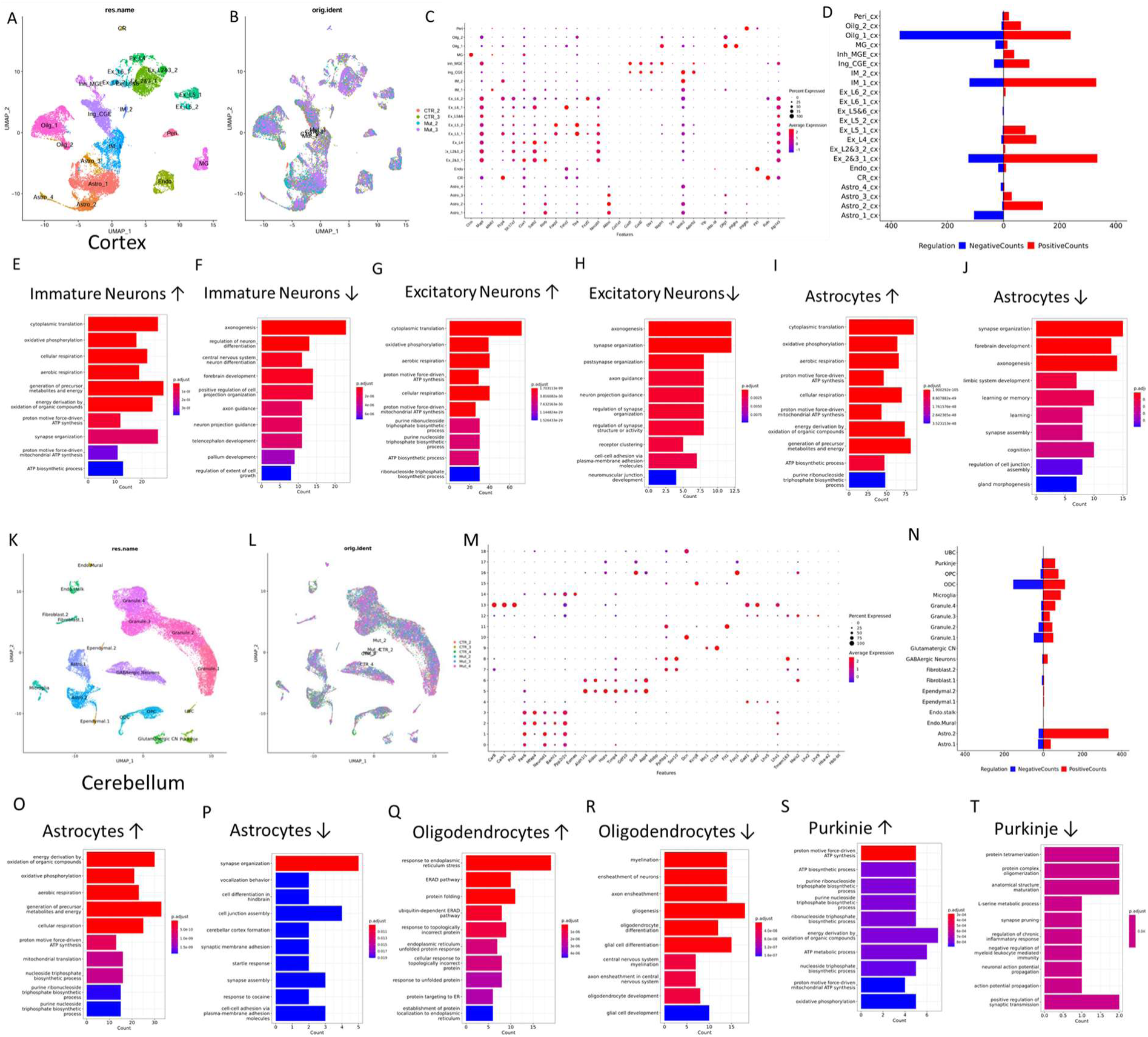
Single-cell transcriptomic alterations in *Pmm2* eKO brain. (A-J) Cortex, (K-T) Cerebellum, UMAP-defined clusters colored by (A) cell population and (B) sample in P7 cortex, (C) Representative marker genes used to define cell identify of UMAP cell population clusters, (D) overview of differentially expressed genes (DEGs) in each cortical cell type, Gene Ontology analysis of DEGs (E) upregulated in immature neurons, (F) downregulated in immature neurons, (G) upregulated in excitatory neurons, (H) downregulated in excitatory neurons, (I) upregulated in astrocytes, (J) downregulated in astrocytes, UMAP-defined clusters colored by (K) cell population and (L) sample in P7 cerebellum, (M) Representative marker genes used to define cell identify of UMAP cell population clusters, (N) overview of differentially expressed genes (DEGs) in each cerebellar cell type, Gene Ontology analysis of DEGs (O) upregulated in astrocytes, (P) downregulated in astrocytes, (Q) upregulated in oligodendrocytes, (R) downregulated in oligodendrocytes, (S) upregulated in Purkinje cells, (T) downregulated in Purkinje cells. *P* indicated via heat map.

### Proteomics and N-glycoproteomics of eKO mice

Given that glycosylation is a post-translational modification and the pathophysiology of PMM2-CDG has generally been assumed to be disruption in protein functions due to hypoglycosylation, we performed proteomic and glycoproteomic analysis of cortex and cerebellum from eKO mice at P12.

In glycoproteomics analysis, we identified and quantified 5,336 N-glycopeptides with 477 unique glycan compositions across 957 N-glycosites spanning 608 glycoproteins. Consistent with glycomics results, we observed aberrant N-glycosylation in both cortex and cerebellum across multiple glycoproteins (**Fig. 9 C and D**). Consistent with the metabolic changes, eKO mice demonstrated more significant hypoglycosylation at the glycopeptide level in cerebellum than in cortex, with an accumulation of glycopeptides bearing smaller oligomannose N-glycans and a decrease in glycopeptides bearing larger oligomannose and complex/hybrid N-glycans. The protein with the highest number of significantly hypoglycosylated glycopeptides was NCAM1. NRCAM and NCAM2 were also significantly hypoglycosylated, among others.

**Fig. 9.**
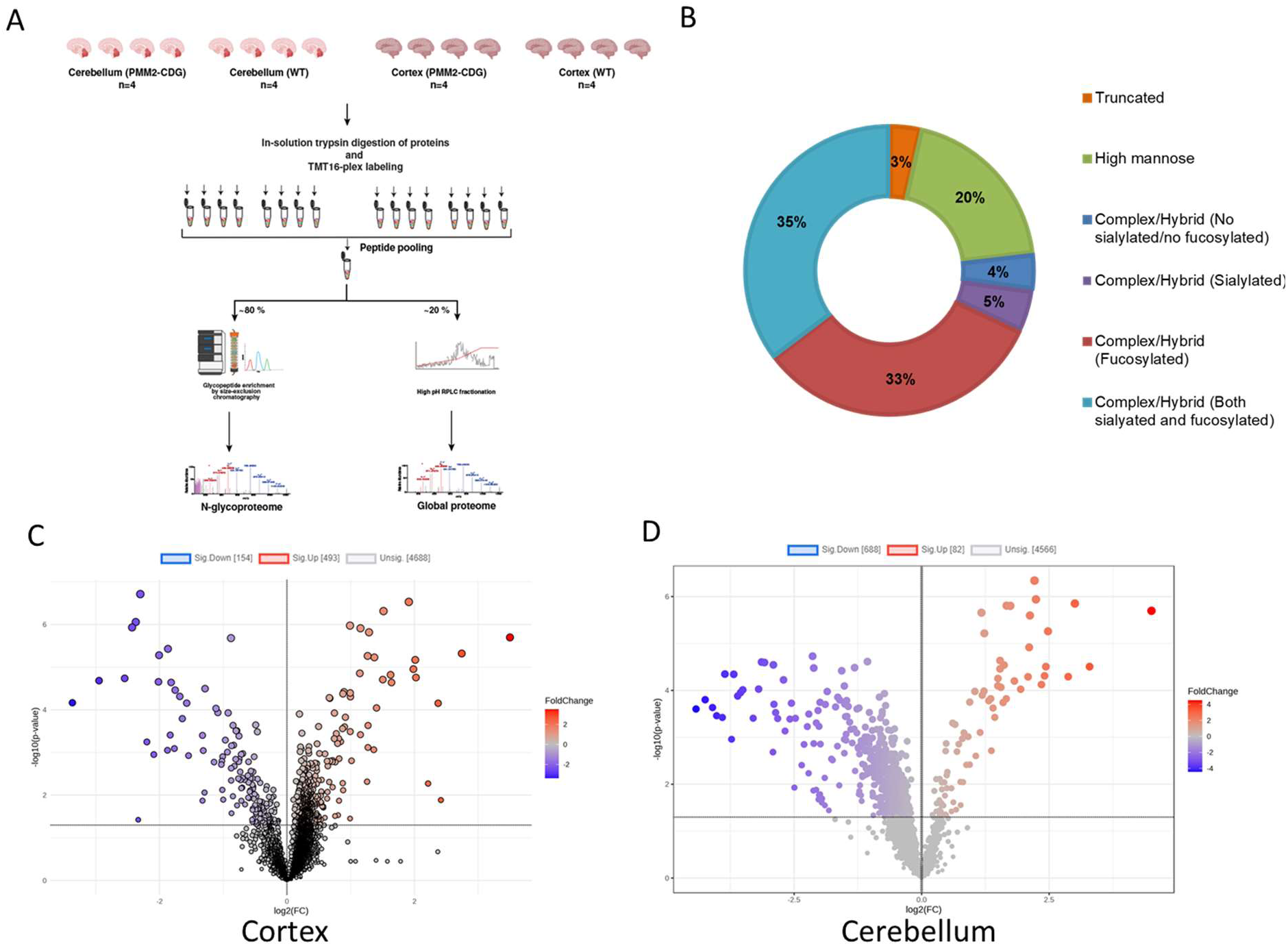
Glycoproteomics analysis in eKO brain. (A) Experimental overview, (B) Glycan types identified in cortex and cerebellum, Volcano plot of glycopeptides in (C) cortex and (D) cerebellum.

### Proteomics and N-glycoproteomics of human PMM2-CDG cerebellum

We previously reported a PMM2-CDG patient who died at 6 months of age due to bleeding complications during attempts to surgically address his pericardial effusion^26^. Clinical autopsy provided opportunity to study the brain of this PMM2-CDG patient with developmental delay, but who had not yet clinically manifest seizures and who died of causes unrelated to his neurologic disease. Autopsy demonstrated cerebellar hypoplasia typical of PMM2-CDG with striking loss of Purkinje cells (**Fig. 10**). The autopsy also provided opportunity to obtain a sample of cerebellum for proteomic and glycoproteomic analysis (**Fig. 11**). Deidentified control cerebellum samples were obtained through clinical pathology and brain biobank repositories from similarly aged individuals who had died of likely non-genetic causes. Proteomic analysis identified 228,515 peptides from 7,971 proteins. Glycoproteomics analysis of human brain enabled identification and quantification of 2,969 N-glycopeptides with 330 unique glycan compositions across 642 N-glycosites spanning 394 glycoproteins. In terms of glycan subtypes, the percent composition of different glycan classes in the human cerebellum was comparable to the mouse cerebellum (**Fig. 11 B**). PMM2-CDG cerebellum demonstrated hypoglycosylation of many of the same glycoproteins identified as statistically hypoglycosylated in the eKO mouse cerebellum, including NCAM1, NRCAM, and NCAM2, thus providing validation of our eKO mouse model as a model of PMM2-CDG brain pathology. Additionally, hypoglycosylation of ATP1B2, GRM7, P2RX7, CACNA2D2, and GRID2 support disruptions in synaptic transmission as a pathogenic mechanism of PMM2-CDG neurological symptoms.

**Fig. 10.**
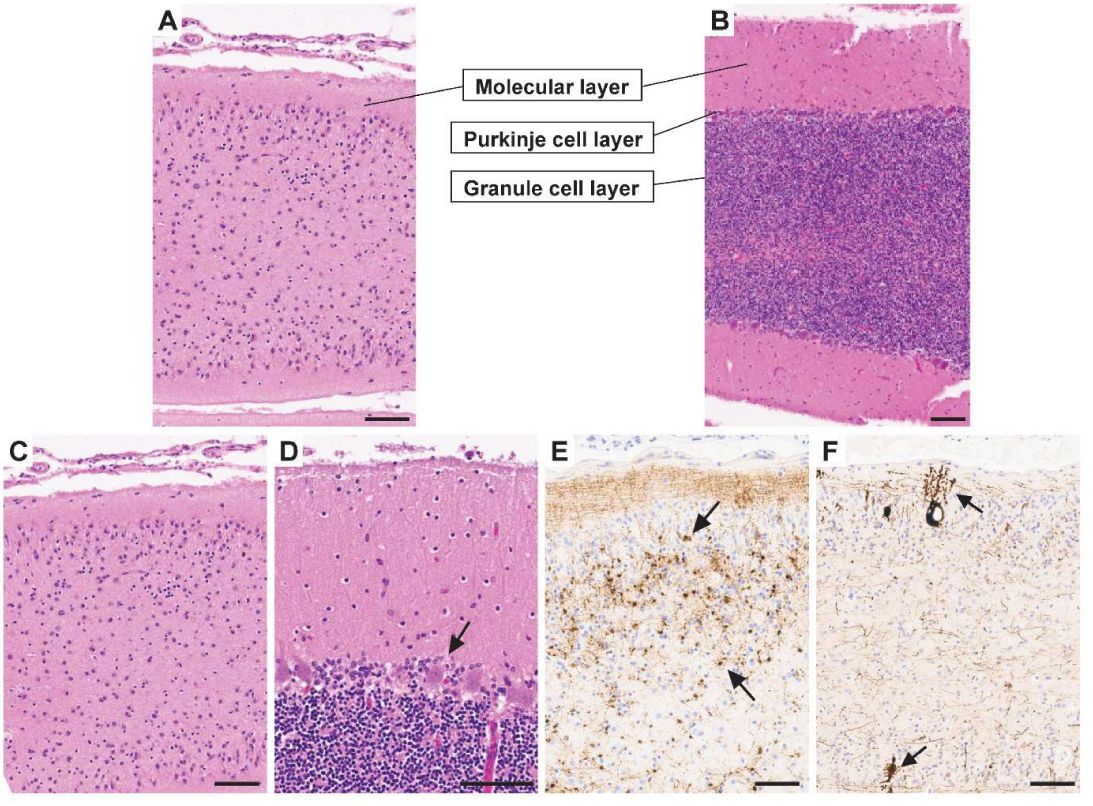
Pathology of PMM2-CDG cerebellum. (A-B) Low-power H&E images show the overall thickness of cerebellar folia from the patient (A) and an age-matched normal counterpart (B). The patient’s specimen exhibits significant folial atrophy. (C) Higher power H&E image from (A, patient) reveals notable architectural disruption in the cerebellar cortex, including loss of Purkinje cells and granule neurons with background gliosis. (D) Higher power H&E image from (B, normal) serves as a comparison delineating the normal lamination of the cerebellar cortex (from top to bottom: molecular layer, Purkinje cell layer (arrow), and granule cell layer). (E) Synaptophysin stain highlights scattered remaining granule cells within the cortex (arrow). (F) Neurofilament stain (NF2F11) demonstrates a marked reduction in axonal processes and highlights rare residual Purkinje cells with dendritic swelling and occasional axonal torpedoes (arrows). Scale bars in all (A-F) equal to 100 μm.

**Fig. 11.**
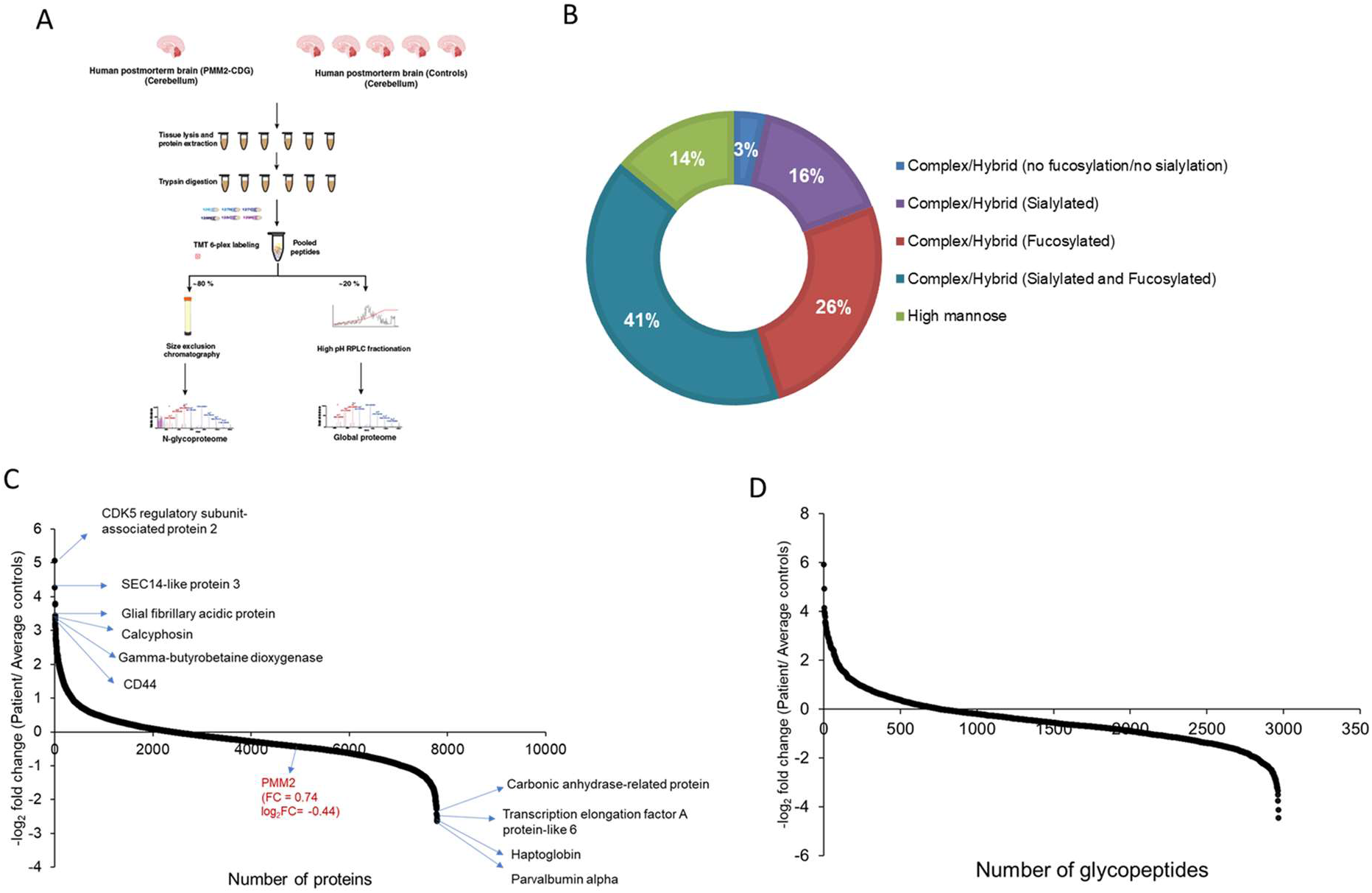
Proteomics and Glycoproteomics analysis in PMM2-CDG cerebellum. (A) Experimental overview, (B) Glycan types identified in human cerebellum, Waterfall plot of (C) peptides and (D) glycopeptides in PMM2-CDG cerebellum.

## Discussion

We report the first mouse model of PMM2-CDG with prominent cerebellar pathology, including ataxia and cerebellar vermian hypoplasia, defining neurological manifestations of PMM2-CDG. To develop a mouse with these defining neurological features required loss of PMM2 during mid-embryonic development in neuronal and glial cell precursors, rather than in post-mitotic neurons or astrocytes, revealing a neurodevelopmental pathogenic mechanism to PMM2-CDG brain pathology. Notably, these was despite similar reductions in bulk brain PMM enzymatic activity in animal models with loss of PMM2 during mid-embryonic development or in post-mitotic neurons.

Other than the vermian hypoplasia, histologic assessment of the brain of affected mice was largely normal, suggesting functional rather than structural abnormalities. Multiomics evaluation identified perturbations throughout the brain with the greatest disruption in the cerebellum, suggesting particular reliance on N-glycosylation in the developing cerebellum, consistent with PMM2-CDG patient manifestations. We observed patterns of dysregulated transcriptional pathways replicated across cell clusters in cortex and cerebellum with transcriptional upregulation of genes related to unfolded protein response and ER stress pathways, findings which were also recently reported in transciptomic analysis of a zebrafish model of PMM2-CDG^10^. We also observed broad transcriptional changes across cell clusters related to oxidative phosphorylation and mitochondrial function, suggestive of mitochondrial dysfunction and impaired glucose metabolism, supportive of prior reports of multiomic analysis of a PMM2-CDG brain organoid model^9^. We also observed that many of the downregulated genes are associated with neurogenesis, neuronal differentiation, axon guidance, synapse organization, and cortical development, also consistent with findings in the human brain organoid model of PMM2-CDG^9^.

The polyol sorbitol, generated by aldose reductase, has been proposed as a toxic metabolite contributor to pathology in PMM2-CDG due to pathogenic metabolic rewiring^11,12^ and aldose reductase inhibitors are in development as novel therapeutics for PMM2-CDG, including a recent phase III clinical trial of epalrestat (NCT04925960). Interestingly, consistent with metabolic findings in PMM2-CDG brain organoids^9^, the polyol pool (which contains sorbitol) was not altered in the brains of our mice, suggesting the influence of sorbitol accumulations on PMM2-CDG disease pathogenesis may be less relevant in the central nervous system. Alternatively, rather than shunting accumulating Man-6-P through sugar alcohol reductase pathways, we observed evidence of increased metabolites in hexosamine biosynthesis and the pentose phosphate pathway. Transcriptional abnormalities in hexosamine biosynthesis pathways were also recently reported in transciptomic analysis of a zebrafish model of PMM2-CDG^10^.

Glycoproteomics identified hypoglycosylation of various neural cell adhesion molecules, synaptic proteins, and cellular signaling receptors. Neural cell adhesion molecules in particular are multifunctional proteins with roles in synaptic pruning and adhesion. Our results suggest abnormal synaptic transmission as a pathogenic mechanism in PMM2-CDG neurological dysfunction. Glycoproteomic disruptions were replicated in post-mortem PMM2-CDG cerebellum, confirming patient-applicability of these findings. These results suggest novel avenues for future research and therapeutic development guided by understanding pathogenic mechanisms of disease in PMM2-CDG.

We highlight ongoing challenges and shortcomings of our mouse model of PMM2-CDG. Ours, and prior mouse models of PMM2-CDG, suggest some organ-system differences between mouse and human. It appears that the degree of reduction of PMM2 enzyme activity to produce neurological pathology exceeds a threshold that is compatible with life in other organs. Producing the core neurological symptoms of PMM2-CDG in our mouse model required a drastic reduction in PMM2 enzyme activity during mid-embryonic development achieved by combining both a catalytically inactive (R137H) *Pmm2* allele and a frameshift loss-of-function *Pmm2* allele. A similar reduction of PMM2 activity later in neurodevelopment failed to produce neurological symptoms, demonstrating a key neurodevelopmental reliance on PMM2, and presumably N-glycosylation, for the neurological symptoms. However, earlier neurodevelopmental stages prior to the expression of Cre experienced normal PMM2 activity, whereas human patients experience hypomorphic PMM2 enzyme activity from conception. However, mouse models with hypomorphic PMM2 enzyme activity from conception have so far failed to recapitulate the neurological aspects of PMM2-CDG, including a compound heterozygous mouse model combining the two most common human pathogenic variants^13^.

A key neurological difference between our mouse model and PMM2-CDG patient brains is that post-mortem pathology analysis of patient brains, including the patient we have included in this study, have demonstrated striking loss of Purkinje cells while Purkinje cells in our eKO mice do not differ in number from control littermates. Autopsy findings in the medical literature have generally been from severely affected PMM2-CDG patients and there is sparse information regarding the status of Purkinje cells in more mildly-affected PMM2-CDG patients. It remains unclear whether PMM2-CDG neurological symptoms result from dysfunctional Purkinje cells, degeneration of Purkinje cells, the failure of Purkinje cells to initially develop, or some combination of the above and how differing levels of PMM2 enzymatic activity may differentially contribute to any of these possibilities. Even across varying degrees of patient severity, the cerebellum is nearly always affected in PMM2-CDG patients and cerebellar dysfunction is their main daily functional limitation^5^. Of note, Purkinje cells are born during the earliest stages of cerebellar neurogenesis and are among the first to be generated in the cerebellar cortex, generally arising between P10-P12.5 in mice^27^, which is earlier than Nestin-Cre is expressed in our model, which is around E12.5^28^. The continued presence of Purkinje cells in our mouse model at numbers similar to control littermates suggests that Purkinje cells degeneration is not a primary driver of pathology, even with severe PMM2 deficiency, and raises questions regarding how much functional improvement can be anticipated from glycosylation restoration therapies under therapeutic development for treating primary CNS manifestations of PMM2-CDG, including gene therapy^29^. Future studies using these *Pmm2* alleles with Cre-lines exhibiting even earlier brain expression so as to impact the earliest stages of cerebellar neurogenesis may provide insight into these questions. Future studies of our nKO mouse model might also be informative regarding ongoing requirements for PMM2 and N-glycosylation in neuronal maintenance and function.

Our pairing of a conditional *Pmm2* mouse allele with a catalytically inactive knock-in allele (R137H) results in a powerful and flexible tool incorporating spatial-temporal control over glycosylation dependent on the presence of Cre recombinase to enable study of organ-specific pathogenic mechanisms of PMM2-CDG and biological functions of N-glycans, as well as generate preclinical models to validate novel therapeutic strategies. It will be a valuable tool for study of other organ systems using different Cre lines, such as the role of PMM2 in the liver as many PMM2-CDG patients also have liver manifestations^30–32^. Pathogenic mechanisms of PMM2-CDG symptoms will vary across organ systems and have variable responses to glycosylation restoration therapeutic approaches. Illustrative of this concept is the observation of prominent renal tubular cysts in our eKO model, which we attribute to ectopic expression of Nestin-Cre in the renal tubule cells of the kidneys. PMM2-CDG patients have a spectrum of renal disease, with cystic kidneys having been reported in some patients^25^. Furthermore, a specific *PMM2* promoter variant (c.-167G>T) appears to uniquely impact binding for the transcription factor ZNF143, important in human kidney and pancreatic β cells^33^, and causes a clinically recognizable hyperinsulinism and polycystic kidney disease (HIPKD) variant form of PMM2-CDG^33–39^ without significant involvement of other organ systems, including the neurological system. PMM2-CDG (HIPKD variant) may represent a naturally-occurring human manifestation of our organ-specific approach and a disease that could benefit from modeling loss of PMM2 in kidney and pancreas cells in a cell-type specific manner.

## Materials and Methods

### Human subjects

This study was performed in accordance with ethical principles for medical research outlined in the Declaration of Helsinki. All relevant approvals from the Children’s Hospital of Philadelphia Institutional Review Board were obtained as well as written informed consent from the patients’ guardians before inclusion in the study. Control samples were obtained post-mortem through the Children’s Hospital of Philadelphia Department of Pathology and as unaffected biopsies of cerebellum from the Children’s Brain Tumor Network biorepository^40^. Control samples were deemed by IRB to not constitute human subjects research as they were provided as deidentified specimens.

### Mouse strains and genotyping

We identified a floxed allele of *Pmm2* generated by the European Conditional Mouse Mutagenesis Program (EUCOMM)^16^, in which exon 3 of *Pmm2* is flanked by two *LoxP* sites cryopreserved at the Canadian Mouse Mutant Repository (CMMR, Toronto), re-derived this mouse line and successfully established a homozygous line of floxed *Pmm2* (*Pmm2*^fl/fl^) in C57Bl/6J background. We obtained the R137H knock-in mouse line (*Pmm2*^R137H/+^, JAX Stock No. 031897)^13^ and Cre lines (Snap25-IRES2-Cre, *Snap25*^Cre/+^, JAX Stock No. 023525^17,18^; B6.Cg-Tg(Gfap-cre)77.6Mvs/2J, Tg(Gfap-Cre), JAX Stock #: 024098^19^; B6.Cg-Tg(Nes-cre)1Kln/J, nestin-Cre, JAX Stock #: 003771^20^) from Jackson Labs. All lines were maintained in the C57BL/6J background. All progeny were genotyped with qPCR by Transnetyx, Inc (Cordova, TN).

### Breeding

Given that the Pmm2^R137H/+^ allele generates a stable transcript and catalytically inactive protein, we initially crossed the *Pmm2*^fl^ allele with the different Cre lines and then bred resulting mice carrying both *Pmm2^fl^*^/+^ and Cre to *Pmm2*^fl/fl^ mice in order to generate mice for molecular validation studies. Subsequently, to prevent germline recombination of the *Pmm2*^fl^ allele from ectopic germline Cre expression, we crossed *Pmm2*^R137H/+^ mice with the different Cre lines and bred resulting mice carrying both *Pmm2*^R137H/+^ and Cre to *Pmm2*^fl/fl^ mice in order to generate compound heterozygote mice where one allele of *Pmm2* is floxed and one allele bears R137H point mutation (*Pmm2*^R137H/fl^) for experimental studies.

### Animal Husbandry

Experiments were conducted in accordance with the ethical guidelines of the National Institutes of Health and with the approval of the Institutional Animal Care and Use Committees of the University of Pennsylvania and Children’s Hospital of Philadelphia. Mice were group housed in cages of 2 to 5 on a 12-hour light/12-hour dark cycle with food and water provided ad libitum. Male littermates and female littermates were weaned at 3-4 weeks of age and housed in sex-segregated, mixed genotype cages.

### Molecular validation

Genomic DNA and RNA was isolated from mouse cortex using the AllPrep DNA/RNA purification kit (Qiagen, Germantown, MD). Genomic recombination assay was performed using PCR primers that flank exon 3 of mouse *Pmm2* (F: 5’-TGAGAGCTCACCTGGCATCATCCATC-3’ R: 5’-ACCTGGGCCTCTTTCACAGCTCCTC-3’) and EmeraldAmp PCR Master Mix (Takara Bio, San Jose, CA). PCR cycling parameters were as follows: 1 cycle of 95°C for 30 seconds; 35 cycles of 95°C for 30 seconds, 60°C for 30 seconds, 72°C for 2.5 min; and 1 cycle of 72°C for 10 min. cDNA was made from isolated RNA using the High Capacity cDNA Reverse Transcription Kit (Applied Biosystems, Foster City, CA) according to the manufacturer’s instructions. TaqMan gene expression assays (Applied Biosystems, Foster City, CA) were used to detect mouse *Pmm2* exon 2 to 3 junction (Mm00450349_m1) and *Hprt* (Mm03024075_m1). Reactions contained 0.5 µL 20x assay mix, 5 µL 2x TaqMan reaction mix, and 1 µL cDNA and were performed in triplicate. Quantitative RT-PCR was performed on an Applied Biosystems 7900HT real-time PCR system. Relative expression differences were calculated by the delta delta Ct method using *Hprt* as the housekeeping gene. Western blots were performed following standard protocols. Samples were separated on Novex NuPAGE 4-12% Bis Tris Protein Gels (Invitrogen) using SDS-PAGE gel electrophoresis and Invitrogen Novex NuPAGE MOPS SDS Running Buffer (Cat. No NP0001, Waltham, MA). Samples were then transferred onto PVDF membanes (Cat. No. GE10600002, Whatman, Little Chalfont We used rabbit anti-PMM2 antibody (orb412234, Biorbyt) and mouse anti-GAPDH antibody (Cat. No. MA5-15738, Thermo Scientific, Waltham, MA) followed by IR labelled anti-rabbit or anti-mouse secondary antibody (Cat.No. 926-68020, Thermo Scientific, Waltham, MA). Membranes were visualized on a Li-Cor imager (Odyssey, Lincoln, NE), and densitometric assessment was performed using Image Studio (Li-Cor).

### PMM enzymatic activity

PMM enzyme activity was measured in brain samples by established PMM enzymatic activity assay^23^ providing Man-1-P as a substrate for PMM2 and subsequently fluorometrically measuring the difference in generated nicotinamide adenine dinucleotide phosphate (NADPH). Briefly, cortex and cerebellum were collected, rinsed in ice cold PBS, and flash frozen in liquid nitrogen. Samples were thawed on ice, homogenized in homogenate buffer (25 mmol/L HEPES buffer - pH 7.1, 25 mmol/L KCl, 0.02% (w/v) Na-azide) using a rotary pestle and sonication, and frozen at −80 ⁰C overnight to lyse the tissue. Total protein was determined by a Bradford protein assay (BioRad) and adjusted to a concentration of 2.5-3.0 µg/uL with additional homogenate buffer. 50 µl of each lysate was combined with 190 µl of PMM reaction mixture (Homogenate buffer, 0.8 mmol/L NADP, 0.1% Inactivated BSA, Man-1,6-bisphosphate, 0.3 mmol/L DTT, and intermediary enzymes [mannose-6 phosphate isomerase (MPI), phosphoglucose isomerase (PGI), glucose-6-phosphate dehydrogenase (G6PD)]). Man-1-Phosphate was added at time 0 to achieve a concentration of 0.7 mmol/L. Absorbance was read at 340 nm at 30 min and 40 min by a Synergy H1 plate reader (BioTek) at 37 ⁰C and PMM activity was determined by the change in absorbance and calculated in nmol/h/mg of total protein.

### Growth and Clinical Veterinary Necropsy Evaluation

Mice were weighed weekly. Complete clinical veterinary necropsy was performed by University of Pennsylvania Comparative Pathology Core collecting organ weights, formalin-fixing and paraffin embedding organs, sectioning, and staining with hematoxylin and eosin following standard clinical procedures. The pathological assessment was performed by trained veterinary pathologists in a blinded fashion without knowledge of the experimental group distribution and genotype of the animals.

### Immunohistologic evaluation of brain

Mice were sacrificed at postnatal days P10 and P17 and underwent transcardial perfusion with PBS followed by 4% paraformaldehyde. Brains were extracted from the skull and post-fixed in 4% paraformaldehyde overnight. Collected brains were embedded in paraffin before microtome sectioning at a thickness of 7 μm. For immunohistochemistry, cortex sections were rehydrated in a series of 100% xylene and 100%–70% ethanol washes before antigen retrieval in a 10 mM sodium citrate solution. Slides were incubated in primary antibodies Calbindin-D (C9848, Millipore, anti-mouse, 1:500), S100b (ab52642, Abcam, anti-rabbit, 1:500), CTIP2 (ab18465, Abcam, anti-rat, 1:500), Cux1 (11733-1-AP, Proteintech, anti-rabbit, 1:500) and 5% normal goat serum in PBS at 4° C overnight. After PBS washes the next day, slides were incubated in secondary antibodies (Alexa Fluor 488 anti-mouse, Cy3 anti-rabbit), 1:500 Hoechst 33342 (Invitrogen #H21492), and 5% normal goat serum in PBS for 3 h. Slides were then washed and mounted with Fluoromount-G (SouthernBiotech #0100-01).

### Cell counting, Cortical and Cerebellar Size Measurements

For cortical and midsagittal cerebellar measurements (including area and cortex width) at P10 and P17, measurements were made using H&E-stained sections for each animal using ImageJ (FIJI)^41^. Cerebellum surface area measurements were taken from whole brain images using ImageJ (FIJI). Cell counting analyses were completed using Photoshop (Adobe v 22.2.0.). At P10 and P17, all Calbindin-positive cells were counted throughout the entire midsagittal area of the cerebellum.

### Behavioral analysis

Mouse behavioral testing was performed in mice beginning at 10-12 weeks of age. Mice were habituated to handling for two weeks prior to starting behavioral testing. Mice were habituated to the testing room for at least 1 hour before testing (except for context- and cue-dependent fear-conditioning assays), which was always performed at the same time of day. Testing was performed in the same order for each animal, beginning with OFM assay, then the EZM assay, Y-maze, 3-chambered social-approach assay, accelerating rotarod assay, context- and cue-dependent fear conditioning, followed by two weeks recovery, then repetitive behavior assay, overnight nesting, and olfaction. Investigators were blinded to genotype while conducting and scoring behavioral assays.

### In vivo Seizure Monitoring

Video-EEG recordings were collected as previously described^42^ using a wireless EEG system (Data Sciences International, St. Paul, MN) and standard surgical approaches with the alteration that given the pre-weaning mortality phenotype of affected mice, P17-P18 preweanling mice were implanted with EEG leads followed by immediate recording and then sacrificed when recording was completed. Mice were anesthetized with isoflurane and four small burr holes were made in the skull above the mouse motor and barrel cortices. A telemetry device containing four electrode leads was implanted subcutaneously on the back of the mouse. The insulation on the positive and negative leads was removed and the exposed wire was manually bent to create a relatively flat terminal to place on the surface of the dura. The leads were stably secured in the head cap via adhesive cement (C&B-Metabond, Parkell Inc, Brentwood, NY). Once the incision was sutured, the mice were given local treatment with antibiotic ointment (OTC Generics, Patterson Veterinary, Houston, TX). Continuous video and EEG recordings were then collected over a brief recording period of up to 8.5 hours using the Ponemah Software System (DSI, St. Paul, MN) and the EEG signal was acquired at 500 Hz. The aligned video and EEG signals are accessed using Neuroscore (DSI) software. First, the EEG signals are preprocessed by filtering with a powerline filter (60 Hz notch filter) followed by 1Hz high pass filtering.

Then, an analysis protocol is designed in the Neuroscore software for spike and spike train detection to determine periods of abnormal EEG. Spikes were detected when the EEG signal had a minimum amplitude of 200 µV and was greater than the root-mean-squared value of the activity within the preceding minute. A spike train was detected when at least five spikes were detected, the inter-spike interval was between 0.05 and 0.6 seconds, and the duration of the train was at least three seconds. Detected spikes and spike trains were then analyzed manually alongside the recorded video for behavior to classify events into runs of epileptiform spikes or behavioral seizures. Runs of epileptiform spikes were detected as spike-trains that lacked clear behavioral manifestation. Behavioral seizures were detected as spike trains of at least three seconds coincident with mouse loss of balance and/or convulsive movements. All EEG data along with identified events were exported into MATLAB to produce raster plots.

### Sample preparation for metabolomics

Briefly, 5 samples of cortex and cerebellum were collected for each genotype and flash frozen without rinsing in PBS to avoid introduction of phosphate as an interfering substance of phosphorylated metabolites. Samples were kept in −80°C prior to metabolomics experiments. Next, the metabolites were extracted using two phase extraction protocol. Briefly, the brain regions were transferred to lysing matrix tube and 350 μL of ice-cold extraction buffer (80% MeOH, supplemented with internal standard) was added to the sample. Next, the brain regions were lyzed with ribolyzer, the lysate transferred to 1.5 mL Eppendorf tube and placed overnight at −80°C. Further, the samples were centrifuged at 15,000 rpm, 4°C, 20 min and 100 μL of supernatant was transferred to a fresh Eppendorf tube. Then, 35 μL of ddH20 was added, followed by 800 μL of 100% chloroform. The samples were then vortexed and stored at 4°C overnight. After the separation of polar and nonpolar phase, the polar phase was then analyzed by Liquid Chromatography/Mass Spectrometry (LC/MS) (see below).

*Metabolite measurement by LC/MS.* As previously described^12^, extracted metabolites were analyzed by LC/MS. First, 10 μL of sample was separated on a C18 ion-pairing liquid chromatography column and the metabolites further resolved by using Thermo Fisher Q-Exactive Hybrid Quadrupole Orbitrap MS in negative ion mode (full scan 70–1050 m/z, resolution 140,000 at 200 m/z, AGC at 3e6, 512 ms ion fill time). The following ESI settings were used: 50 sheet gas flow rate, spray voltage of 4 kV, auxiliary gas flow rate 15, S-lens RF level of 60, and the capillary temperature at 350°C. The identification of the metabolites was performed in accordance with their m/z ratio and elution time using in-house metabolite standard library and El-Maven v0.12.0/Polly ™Labeled LC-MS Workflow. Finally, metabolite abundances were normalized to the weight of brain region and internal standards. Relative values were established using unaffected mice as reference. Absolute quantification was not performed.

### N-glycomics

Brain regions that had been collected, rinsed in ice cold PBS, and flash frozen were lysed in MS-grade water using a rotary pestle followed by sonication. N-glycans were released from brain lysate using a rapid PNGaseF digestion, derivatized and purified using Rapifluor TM N-glycan preparation kit as described previously^43^. Briefly, an N-hydroxysuccinimide carbamate tag with a modified quinolone is added to the transient glycosylamide group at the reducing end of the glycans. The derivatized N-glycans were purified using a 96 well HILIC plate and analyzed using a flow-injection-ESI-QTOF Mass Spectrometry method. A glycopeptide standard with isotope labelled disialoglycan was used as the internal standard. The abundance of each glycan is reported as % total glycans.

### Single cell Isolation

For cortex and cerebellum tissues collected from P7 mice, animals were euthanized by decapitation and dissections were carried out on an ice-cold surface. Samples were finely minced using a clean blade and maintained in ice-cold DPBS containing 0.01% BSA and 30 μM actinomycin D (ActD) to inhibit transcription, until all samples were processed. Tissue dissociation was performed using the Worthington Papain Dissociation System with a 45-minute enzymatic digestion step, following the manufacturer’s instructions with modification of 15 μM ActD being added to both the papain/DNase digestion buffer and the albumin-ovomucoid inhibitor solution used for density gradient separation.

Following dissociation, cells were pelleted by centrifugation at 100 × g for 6 minutes, then resuspended in 0.5 mL DPBS containing 0.01% BSA (without ActD). Cell numbers were quantified using the Countess II automated counter (Invitrogen). The suspension was split into two 1.5 mL LoBind tubes (Eppendorf), and ice-cold methanol was added dropwise to reach a final concentration of 80% methanol in DPBS. After incubating on ice for 1 hour, fixed cells were stored at –80°C for up to one month.

### Library preparation & sequencing

Gene expression libraries were generated using the Chromium Single Cell 3’ Library Kit v3 (10x Genomics, #1000078), following the manufacturer’s protocol. In brief, 10,000 nuclei per sample were used for the initial transposition reaction and subsequently loaded onto the Chromium Controller, aiming to recover approximately 8,000 nuclei per sample. The resulting barcoded cDNA was then processed to construct gene expression libraries according to standard procedures. Library quality was assessed throughout using an Agilent Bioanalyzer and Thermo Fisher Qubit fluorometer. Sequencing was performed on the NovaSeq 6000 platform (Illumina) with the following cycle configuration: 28 cycles for Read 1, 10 cycles each for i7 and i5 indices, and 90 cycles for Read 2. *Preprocessing of scRNAseq data.* Demultiplexed FASTQ files were aligned to the mouse reference genome (mm10, Gencode release vM13) using Cell Ranger (10X Genomics, version 7.1.0) with default settings. To reduce ambient RNA contamination, we employed CellBender (version 0.3.1)^44^, a probabilistic deep learning-based tool designed to identify and remove background RNA from droplet-based scRNA-seq data. Subsequent transcriptomic analysis was performed using the Seurat R package (version 4.3.0)^45^. For quality control, cells with fewer than 400 or more than 7,000 detected genes, or with mitochondrial gene content exceeding 15%, were excluded. Genes detected in fewer than 10 cells across the dataset were also filtered out. To eliminate potential doublets, Scrublet (version 0.2.3)^46^ was applied, with cells scoring above 0.3 removed from each sample. The dataset was normalized using the NormalizeData function, and highly variable features were identified via FindVariableFeatures. Principal component analysis was performed using RunPCA, and the top 30 principal components were used to compute a shared nearest neighbor graph (FindNeighbors) and generate clusters (FindClusters). Cell type annotation was based on canonical marker genes previously reported in cortex^47^ and cerebellum^48^. Clusters expressing markers from more than two distinct lineages or containing more than 50% predicted doublets were excluded from further analysis. In cortical tissue, cells were broadly classified into glutamatergic neurons, GABAergic neurons, and non-neuronal cell types, such as astrocytes, oligodendrocytes, microglia, and endothelial cells. In the cerebellum, cells were grouped into Purkinje cells, granule neuron lineage cells, unipolar brush cells, and non-neuronal populations, using known marker genes for classification.

### Single-cell RNA-seq statistical analysis

To assess transcriptional changes between *Pmm2* eKO mice and littermate controls and to identify subclass- or cell-type-specific marker genes, we used the FindMarkers function from Seurat. Differentially expressed genes (DEGs) were defined by an adjusted *P* < 0.05. To evaluate enrichment for biological process, clusterProfiler (version 4.10.0)^49^ was used to perform Gene Ontology (GO) enrichment analysis on significantly upregulated and downregulated DEGs in each cell cluster. GO terms with a Benjamini–Hochberg adjusted *P* < 0.05 were considered significantly enriched. *Sample preparation for proteomics and glycoproteomics.* Brain samples were lysed using Bioruptor sonication device in 0.1% DDM (in 100 mM TEABC) supplemented with protease inhibitors. Brain lysates were centrifuged to remove the cell debris. Protein quantities in the obtained supernatant were estimated by BCA assay as per the manufacturer’s instructions. Equal amounts of protein from each sample were first reduced using 10 mM dithiothreitol for 30 min at 55°C on a thermomixer, then alkylated with 40 mM iodoacetamide for 30 min in the dark. The proteins were then digested with trypsin at 37°C overnight with mild shaking on thermomixer. Resulting peptides were desalted using C18 columns and labeled with tandem mass tags (TMT) as per the manufacturer’s protocol and subsequently pooled.

### Fractionation and glycopeptide enrichment

The pooled TMT-labeled peptides were split into two aliquots. One aliquot (∼20% of the total peptides) was resuspended in solvent A (5 mM ammonium formate, pH 9) and fractionated by basic pH reversed phase liquid chromatography (bRPLC) on a reversed phase Waters C18 column (5 μm, 4.6 × 100 mm column) using an increasing gradient of solvent B (5 mM ammonium formate, pH 9, in 90% acetonitrile) on the Ultimate 3000 UHPLC system. Ninety-six fractions were collected and subsequently concatenated into 12 fractions for the proteomics experiment. The other aliquot from the pooled peptides was used to enrich glycopeptides. About 80% of the total peptides was resuspended in 100 μL of 0.1% formic acid and injected into Superdex peptide 10/300 column as described previously.75 The peptides were separated using an isocratic flow of 0.1% formic acid for 130 min and early fractions were collected starting at 10 min after injection. These fractions were subsequently concatenated into 12 fractions for the glycoproteomics experiment.

### Proteome and glycoproteome analysis by liquid chromatography-tandem mass spectrometry (LC-MS/MS)

LC-MS/MS analysis of fractionated samples from both proteomics and glycoproteomics was carried out. The samples were analyzed on an Orbitrap Eclipse mass spectrometer equipped with Ultimate 3000 liquid chromatography system (Thermo Fisher Scientific Inc.). The peptides/glycopeptides were separated on an analytical column (EasySpray 75 μm × 50 cm, C18 2 μm, 100 Å, with a flow rate of 300 nL/min with a linear gradient of solvent B (100% ACN, 0.1% formic acid) over a 155 min gradient.

Precursor ions were acquired at a resolution of 120,000 (at m/z 200) for precursor ions and at a resolution of 30,000 (at m/z 200) for fragment ions. Precursor ions were acquired in the Orbitrap mass analyzer in m/z range of 350-1,700 for proteomics and 375-2,000 for glycoproteomics. The fragmentation was carried out using higher-energy collisional dissociation (HCD) method using normalized collision energy of 35 for proteomics or stepped HCD (15, 25, 40) for glycoproteomics. The scans were acquired in top-speed method with 3 s cycle time between MS and MS/MS.

### Proteomics and glycoproteomics data analysis

The proteomics data were searched using Sequest search engine in Proteome Discoverer 2.5 against the Uniprot Human Reviewed protein sequences (20,432 entries) and the glycoproteomics data using the publicly available software pGlyco version 2.2 with an in-built glycan database containing 8,092 entries. Two missed cleavages were allowed for both proteomics and glycoproteomics analysis. For proteomics data, error tolerance for precursor and fragment ions was set to 10 and 0.02 Da ppm, respectively, and for glycoproteomics data it was set to 10 ppm and 20 ppm, respectively. Cysteine carbamidomethylation, TMT mass on lysines and peptide N-termini were set as fixed modification with oxidation of methionine as a variable modification. False discovery rate (FDR) was set to 1% at the peptide-spectrum matches (PSMs), peptide, protein, and glycopeptides levels. For proteomics, quantitation of peptides across PMM2-CDG and control brain regions was done using TMT reporter ion intensities using “reporter ion quantifier” node. To quantify glycopeptides, reporter ion quantification was performed for glycoproteomics raw files in Proteome Discoverer version 2.5 and glycopeptide IDs obtained from pGlyco 2.2 were matched with quantitation on a scan-to-scan basis (MS/MS). Two-sample student’s t test with unequal group variance was used to identify differentially expressed proteins and glycopeptides in PMM2 deficient hCOs. For mitochondrial proteins, the data were searched against the annotated gene list which was generated from MitoCarta 3.0.83

### Metabolomics data analysis

Metabolomics data were analyzed with the El Maven Polly software using the m/z and retention time (validated by the in-house standards library and internal standards). Metabolite abundances were normalized to brain region weight. Relative metabolite abundances were calculated by comparing eKO samples to littermate control samples. Absolute quantification was not performed. Statistics was performed using GraphPad prism (version 10 for MacBook). Metabolanalyst was used to generate PCA plots and further analyze the data.

### Statistics

For statistical analyses we used Welch’s unpaired, 2-tailed t-test, except as follows: genotype distributions, χ2 test against expected Mendelian distribution; 3-chambered social choice assay and olfaction, 2-way ANOVA with Fisher’s Least Significant Differences test; rotarod, generalized linear mixed effects model with Geisser-Greenhouse correction and Fisher’s Least Significant Differences test; nesting, unpaired Mann-Whitney test; spikes, 1-way ANOVA. All graphs are plotted using Prism (GraphPad). Level of significance was set at *P* ≤ 0.05. In our figures, * is used to denote all 0.01 < *P* < 0.05, ** for 0.001 < *P* < 0.01, *** for 0.0001 < *P* < 0.001, and **** for *P* < 0.0001. Data are represented as mean ± SD, except rotarod where errors bars represent ± SEM and nesting where the line indicates median. For each analysis of mouse brain immunohistochemistry, between three and six animals for each genotype group were assessed. Statistical significance was assessed using an unpaired, two-tailed, student’s t-test using GraphPad Prism (version 9.1.0 for Windows, GraphPad Software, San Diego, CA). Given we only had a single PMM2-CDG sample to compare to 5 control samples, we were unable to perform typical statistical comparison, we therefore calculated average and standard deviation values for peptides and glycopeptides within the control samples and compared with the PMM2-CDG sample and used a cut-off of more than 2 standard deviations from the control mean to determine whether a peptide or glycopeptide were significantly depleted or enriched in the study.

## Acknowledgments

We would like to acknowledge Joshua Ross, Rashmi Yadav, and Siddharth Sobti for their technical contributions to the project, as well as the services of the University of Pennsylvania Comparative Pathology Core in the clinical pathological assessment of mice and the Children’s Hospital of Philadelphia Department of Pathology and the Children’s Brain Tumor Network for providing post-mortem control samples.

## REFERENCES

1. Schollen, E., Kjaergaard, S., Legius, E., Schwartz, M., and Matthijs, G. (2000). Lack of Hardy-Weinberg equilibrium for the most prevalent PMM2 mutation in CDG-Ia (congenital disorders of glycosylation type Ia). European journal of human genetics: EJHG 8, 367–371. 10.1038/sj.ejhg.5200470.

2. Altassan, R., Peanne, R., Jaeken, J., Barone, R., Bidet, M., Borgel, D., Brasil, S., Cassiman, D., Cechova, A., Coman, D., et al. (2019). International clinical guidelines for the management of phosphomannomutase 2-congenital disorders of glycosylation: Diagnosis, treatment and follow up. Journal of inherited metabolic disease 42, 5–28. 10.1002/jimd.12024.

3. Sparks, S.E., and Krasnewich, D.M. (1993). PMM2-CDG (CDG-Ia). In GeneReviews((R)), M.P. Adam, H.H. Ardinger, R.A. Pagon, S.E. Wallace, L.J.H. Bean, K. Stephens, and A. Amemiya, eds.

4. Schiff, M., Roda, C., Monin, M.L., Arion, A., Barth, M., Bednarek, N., Bidet, M., Bloch, C., Boddaert, N., Borgel, D., et al. (2017). Clinical, laboratory and molecular findings and long-term follow-up data in 96 French patients with PMM2-CDG (phosphomannomutase 2-congenital disorder of glycosylation) and review of the literature. Journal of medical genetics 54, 843–851. 10.1136/jmedgenet-2017-104903.

5. Serrano, M., de Diego, V., Muchart, J., Cuadras, D., Felipe, A., Macaya, A., Velazquez, R., Poo, M.P., Fons, C., O’Callaghan, M.M., et al. (2015). Phosphomannomutase deficiency (PMM2-CDG): ataxia and cerebellar assessment. Orphanet journal of rare diseases 10, 138. 10.1186/s13023-015-0358-y.

6. Aronica, E., van Kempen, A.A., van der Heide, M., Poll-The, B.T., van Slooten, H.J., Troost, D., and Rozemuller-Kwakkel, J.M. (2005). Congenital disorder of glycosylation type Ia: a clinicopathological report of a newborn infant with cerebellar pathology. Acta neuropathologica 109, 433–442. 10.1007/s00401-004-0975-3.

7. Antoun, H., Villeneuve, N., Gelot, A., Panisset, S., and Adamsbaum, C. (1999). Cerebellar atrophy: an important feature of carbohydrate deficient glycoprotein syndrome type 1. Pediatr Radiol 29, 194–198. 10.1007/s002470050571.

8. Ng, B.G., and Freeze, H.H. (2018). Perspectives on Glycosylation and Its Congenital Disorders. Trends Genet 34, 466–476. 10.1016/j.tig.2018.03.002.

9. Radenkovic, S., Budhraja, R., Klein-Gunnewiek, T., King, A.T., Bhatia, T.N., Ligezka, A.N., Driesen, K., Shah, R., Ghesquiere, B., Pandey, A., et al. (2024). Neural and metabolic dysregulation in PMM2-deficient human in vitro neural models. Cell Rep 43, 113883. 10.1016/j.celrep.2024.113883.

10. Matheny-Rabun, C., Mokashi, S.S., Radenkovic, S., Wiggins, K., Dukes-Rimsky, L., Angel, P., Ghesquiere, B., Kozicz, T., Steet, R., Morava, E., and Flanagan-Steet, H. (2024). O-GlcNAcylation modulates expression and abundance of N-glycosylation machinery in an inherited glycosylation disorder. Cell Rep 43, 114976. 10.1016/j.celrep.2024.114976.

11. Ligezka, A.N., Radenkovic, S., Saraswat, M., Garapati, K., Ranatunga, W., Krzysciak, W., Yanaihara, H., Preston, G., Brucker, W., McGovern, R.M., et al. (2021). Sorbitol Is a Severity Biomarker for PMM2-CDG with Therapeutic Implications. Ann Neurol 90, 887–900. 10.1002/ana.26245.

12. Radenkovic, S., Ligezka, A.N., Mokashi, S.S., Driesen, K., Dukes-Rimsky, L., Preston, G., Owuocha, L.F., Sabbagh, L., Mousa, J., Lam, C., et al. (2023). Tracer metabolomics reveals the role of aldose reductase in glycosylation. Cell Rep Med 4, 101056. 10.1016/j.xcrm.2023.101056.

13. Chan, B., Clasquin, M., Smolen, G.A., Histen, G., Powe, J., Chen, Y., Lin, Z., Lu, C., Liu, Y., Cang, Y., et al. (2016). A mouse model of a human congenital disorder of glycosylation caused by loss of PMM2. Human molecular genetics 25, 2182–2193. 10.1093/hmg/ddw085.

14. Thiel, C., Lubke, T., Matthijs, G., von Figura, K., and Korner, C. (2006). Targeted disruption of the mouse phosphomannomutase 2 gene causes early embryonic lethality. Molecular and cellular biology 26, 5615–5620. 10.1128/MCB.02391-05.

15. Schneider, A., Thiel, C., Rindermann, J., DeRossi, C., Popovici, D., Hoffmann, G.F., Grone, H.J., and Korner, C. (2011). Successful prenatal mannose treatment for congenital disorder of glycosylation-Ia in mice. Nature medicine 18, 71–73. 10.1038/nm.2548.

16. Friedel, R.H., Seisenberger, C., Kaloff, C., and Wurst, W. (2007). EUCOMM--the European conditional mouse mutagenesis program. Brief Funct Genomic Proteomic 6, 180–185. 10.1093/bfgp/elm022.

17. Harris, J.A., Hirokawa, K.E., Sorensen, S.A., Gu, H., Mills, M., Ng, L.L., Bohn, P., Mortrud, M., Ouellette, B., Kidney, J., et al. (2014). Anatomical characterization of Cre driver mice for neural circuit mapping and manipulation. Frontiers in neural circuits 8, 76. 10.3389/fncir.2014.00076.

18. Paton, K.M., Selfridge, J., Guy, J., and Bird, A. (2022). Comparative analysis of potential broad-spectrum neuronal Cre drivers. Wellcome Open Res 7, 185. 10.12688/wellcomeopenres.17965.1.

19. Gregorian, C., Nakashima, J., Le Belle, J., Ohab, J., Kim, R., Liu, A., Smith, K.B., Groszer, M., Garcia, A.D., Sofroniew, M.V., et al. (2009). Pten deletion in adult neural stem/progenitor cells enhances constitutive neurogenesis. The Journal of neuroscience: the official journal of the Society for Neuroscience 29, 1874–1886. 10.1523/JNEUROSCI.3095-08.2009.

20. Tronche, F., Kellendonk, C., Kretz, O., Gass, P., Anlag, K., Orban, P.C., Bock, R., Klein, R., and Schutz, G. (1999). Disruption of the glucocorticoid receptor gene in the nervous system results in reduced anxiety. Nature genetics 23, 99–103. 10.1038/12703.

21. Cacheiro, P., Haendel, M.A., Smedley, D., International Mouse Phenotyping, C., and the Monarch, I. (2019). New models for human disease from the International Mouse Phenotyping Consortium. Mammalian genome: official journal of the International Mammalian Genome Society 30, 143–150. 10.1007/s00335-019-09804-5.

22. Luo, L., Ambrozkiewicz, M.C., Benseler, F., Chen, C., Dumontier, E., Falkner, S., Furlanis, E., Gomez, A.M., Hoshina, N., Huang, W.H., et al. (2020). Optimizing Nervous System-Specific Gene Targeting with Cre Driver Lines: Prevalence of Germline Recombination and Influencing Factors. Neuron 106, 37–65 e35. 10.1016/j.neuron.2020.01.008.

23. Van Schaftingen, E., and Jaeken, J. (1995). Phosphomannomutase deficiency is a cause of carbohydrate-deficient glycoprotein syndrome type I. FEBS letters 377, 318–320. 10.1016/0014-5793(95)01357-1.

24. Pirard, M., Achouri, Y., Collet, J.F., Schollen, E., Matthijs, G., and Van Schaftingen, E. (1999). Kinetic properties and tissular distribution of mammalian phosphomannomutase isozymes. The Biochemical journal 339 (Pt 1), 201–207.

25. Altassan, R., Witters, P., Saifudeen, Z., Quelhas, D., Jaeken, J., Levtchenko, E., Cassiman, D., and Morava, E. (2018). Renal involvement in PMM2-CDG, a mini-review. Molecular genetics and metabolism 123, 292–296. 10.1016/j.ymgme.2017.11.012.

26. Hong, X., Alharbi, H., Albokhari, D., Edmondson, A.C., and He, M. (2022). A 6-Month-Old Infant with Severe Failure to Thrive during COVID-19 Pandemic. Clinical chemistry 68, 987–989. 10.1093/clinchem/hvac012.

27. Marti-Clua, J. (2022). Times of neuron origin and neurogenetic gradients in mice Purkinje cells and deep cerebellar nuclei neurons during the development of the cerebellum. A review. Tissue & cell 78, 101897. 10.1016/j.tice.2022.101897.

28. Liang, H., Hippenmeyer, S., and Ghashghaei, H.T. (2012). A Nestin-cre transgenic mouse is insufficient for recombination in early embryonic neural progenitors. Biol Open 1, 1200–1203. 10.1242/bio.20122287.

29. Zhong, M.L., and Lai, K. (2025). AAV-based gene replacement therapy prevents and halts manifestation of abnormal neurological phenotypes in a novel mouse model of PMM2-CDG. Gene Ther 32, 246–254. 10.1038/s41434-025-00525-w.

30. Starosta, R.T., Boyer, S., Tahata, S., Raymond, K., Lee, H.E., Wolfe, L.A., Lam, C., Edmondson, A.C., Schwartz, I.V.D., and Morava, E. (2021). Liver manifestations in a cohort of 39 patients with congenital disorders of glycosylation: pin-pointing the characteristics of liver injury and proposing recommendations for follow-up. Orphanet journal of rare diseases 16, 20. 10.1186/s13023-020-01630-2.

31. Lam, C., Scaglia, F., Berry, G.T., Larson, A., Sarafoglou, K., Andersson, H.C., Sklirou, E., Tan, Q.K.G., Starosta, R.T., Sadek, M., et al. (2024). Frontiers in congenital disorders of glycosylation consortium, a cross-sectional study report at year 5 of 280 individuals in the natural history cohort. Molecular genetics and metabolism 142, 108509. 10.1016/j.ymgme.2024.108509.

32. Johnsen, C., and Edmondson, A.C. (2021). Manifestations and Management of Hepatic Dysfunction in Congenital Disorders of Glycosylation. Clin Liver Dis (Hoboken) 18, 54–66. 10.1002/cld.1105.

33. Cabezas, O.R., Flanagan, S.E., Stanescu, H., Garcia-Martinez, E., Caswell, R., Lango-Allen, H., Anton-Gamero, M., Argente, J., Bussell, A.M., Brandli, A., et al. (2017). Polycystic Kidney Disease with Hyperinsulinemic Hypoglycemia Caused by a Promoter Mutation in Phosphomannomutase 2. Journal of the American Society of Nephrology: JASN 28, 2529–2539. 10.1681/ASN.2016121312.

34. Carney, E.F. (2017). Polycystic kidney disease: PMM2 mutation causes PKD and hyperinsulinism. Nat Rev Nephrol 13, 321. 10.1038/nrneph.2017.58.

35. Moreno Macian, F., De Mingo Alemany, C., Leon Carinena, S., Ortega Lopez, P., Rausell Felix, D., Aparisi Navarro, M., Martinez Matilla, M., Cardona Gay, C., Martinez Castellano, F., and Albiach Mesado, V. (2020). Mutations in PMM2 gene in four unrelated Spanish families with polycystic kidney disease and hyperinsulinemic hypoglycemia. J Pediatr Endocrinol Metab 33, 1283–1288. 10.1515/jpem-2020-0168.

36. Soares, A.R., Figueiredo, C.M., Quelhas, D., Silva, E.S., Freitas, J., Oliveira, M.J., Faria, S., Fortuna, A.M., and Borges, T. (2020). Hyperinsulinaemic Hypoglycaemia and Polycystic Kidney Disease - A Rare Case Concerning PMM2 Gene Pleiotropy. Eur Endocrinol 16, 66–68. 10.17925/EE.2020.16.1.66.

37. Dorval, G., Jeanpierre, C., Moriniere, V., Tournant, C., Bessieres, B., Attie-Bittach, T., Amiel, J., Spaggari, E., Ville, Y., Merieau, E., et al. (2021). Cystic kidney diseases associated with mutations in phosphomannomutase 2 promotor: a large spectrum of phenotypes. Pediatr Nephrol 36, 2361–2369. 10.1007/s00467-021-04953-9.

38. Islam, S., Tekman, M., Flanagan, S.E., Guay-Woodford, L., Hussain, K., Ellard, S., Kleta, R., Bockenhauer, D., Stanescu, H., and Iancu, D. (2021). Founder mutation in the PMM2 promotor causes hyperinsulinemic hypoglycaemia/polycystic kidney disease (HIPKD). Molecular genetics & genomic medicine 9, e1674. 10.1002/mgg3.1674.

39. Kiparissi, F., Dastamani, A., Palm, L., Azabdaftari, A., Campos, L., Gaynor, E., Grunewald, S., Uhlig, H.H., Kleta, R., Bockenhauer, D., and Jones, K.D.J. (2023). Phosphomannomutase 2 (PMM2) variants leading to hyperinsulinism-polycystic kidney disease are associated with early-onset inflammatory bowel disease and gastric antral foveolar hyperplasia. Hum Genet 142, 697–704. 10.1007/s00439-023-02523-7.

40. Lilly, J.V., Rokita, J.L., Mason, J.L., Patton, T., Stefankiewiz, S., Higgins, D., Trooskin, G., Larouci, C.A., Arya, K., Appert, E., et al. (2023). The children’s brain tumor network (CBTN) - Accelerating research in pediatric central nervous system tumors through collaboration and open science. Neoplasia 35, 100846. 10.1016/j.neo.2022.100846.

41. Schindelin, J., Arganda-Carreras, I., Frise, E., Kaynig, V., Longair, M., Pietzsch, T., Preibisch, S., Rueden, C., Saalfeld, S., Schmid, B., et al. (2012). Fiji: an open-source platform for biological-image analysis. Nat Methods 9, 676-682. 10.1038/nmeth.2019.

42. Feng, H., Clatot, J., Kaneko, K., Flores-Mendez, M., Wengert, E.R., Koutcher, C., Hoddeson, E., Lopez, E., Lee, D., Arias, L., et al. (2024). Targeted therapy improves cellular dysfunction, ataxia, and seizure susceptibility in a model of a progressive myoclonus epilepsy. Cell Rep Med 5, 101389. 10.1016/j.xcrm.2023.101389.

43. Chen, J., Li, X., Edmondson, A., Meyers, G.D., Izumi, K., Ackermann, A.M., Morava, E., Ficicioglu, C., Bennett, M.J., and He, M. (2019). Increased Clinical Sensitivity and Specificity of Plasma Protein N-Glycan Profiling for Diagnosing Congenital Disorders of Glycosylation by Use of Flow Injection-Electrospray Ionization-Quadrupole Time-of-Flight Mass Spectrometry. Clinical chemistry 65, 653–663. 10.1373/clinchem.2018.296780.

44. Fleming, S.J., Chaffin, M.D., Arduini, A., Akkad, A.D., Banks, E., Marioni, J.C., Philippakis, A.A., Ellinor, P.T., and Babadi, M. (2023). Unsupervised removal of systematic background noise from droplet-based single-cell experiments using CellBender. Nat Methods 20, 1323–1335. 10.1038/s41592-023-01943-7.

45. Stuart, T., Butler, A., Hoffman, P., Hafemeister, C., Papalexi, E., Mauck, W.M., 3rd, Hao, Y., Stoeckius, M., Smibert, P., and Satija, R. (2019). Comprehensive Integration of Single-Cell Data. Cell 177, 1888-1902 e1821. 10.1016/j.cell.2019.05.031.

46. Wolock, S.L., Lopez, R., and Klein, A.M. (2019). Scrublet: Computational Identification of Cell Doublets in Single-Cell Transcriptomic Data. Cell Syst 8, 281–291 e289. 10.1016/j.cels.2018.11.005.

47. Yao, Z., Liu, H., Xie, F., Fischer, S., Adkins, R.S., Aldridge, A.I., Ament, S.A., Bartlett, A., Behrens, M.M., Van den Berge, K., et al. (2021). A transcriptomic and epigenomic cell atlas of the mouse primary motor cortex. Nature 598, 103–110. 10.1038/s41586-021-03500-8.

48. Kozareva, V., Martin, C., Osorno, T., Rudolph, S., Guo, C., Vanderburg, C., Nadaf, N., Regev, A., Regehr, W.G., and Macosko, E. (2021). A transcriptomic atlas of mouse cerebellar cortex comprehensively defines cell types. Nature 598, 214–219. 10.1038/s41586-021-03220-z.

49. Yu, G., Wang, L.G., Han, Y., and He, Q.Y. (2012). clusterProfiler: an R package for comparing biological themes among gene clusters. OMICS 16, 284–287. 10.1089/omi.2011.0118.

