## Supplemental Figures for "Novel mouse model reveals neurodevelopmental origin of PMM2-CDG brain pathology"

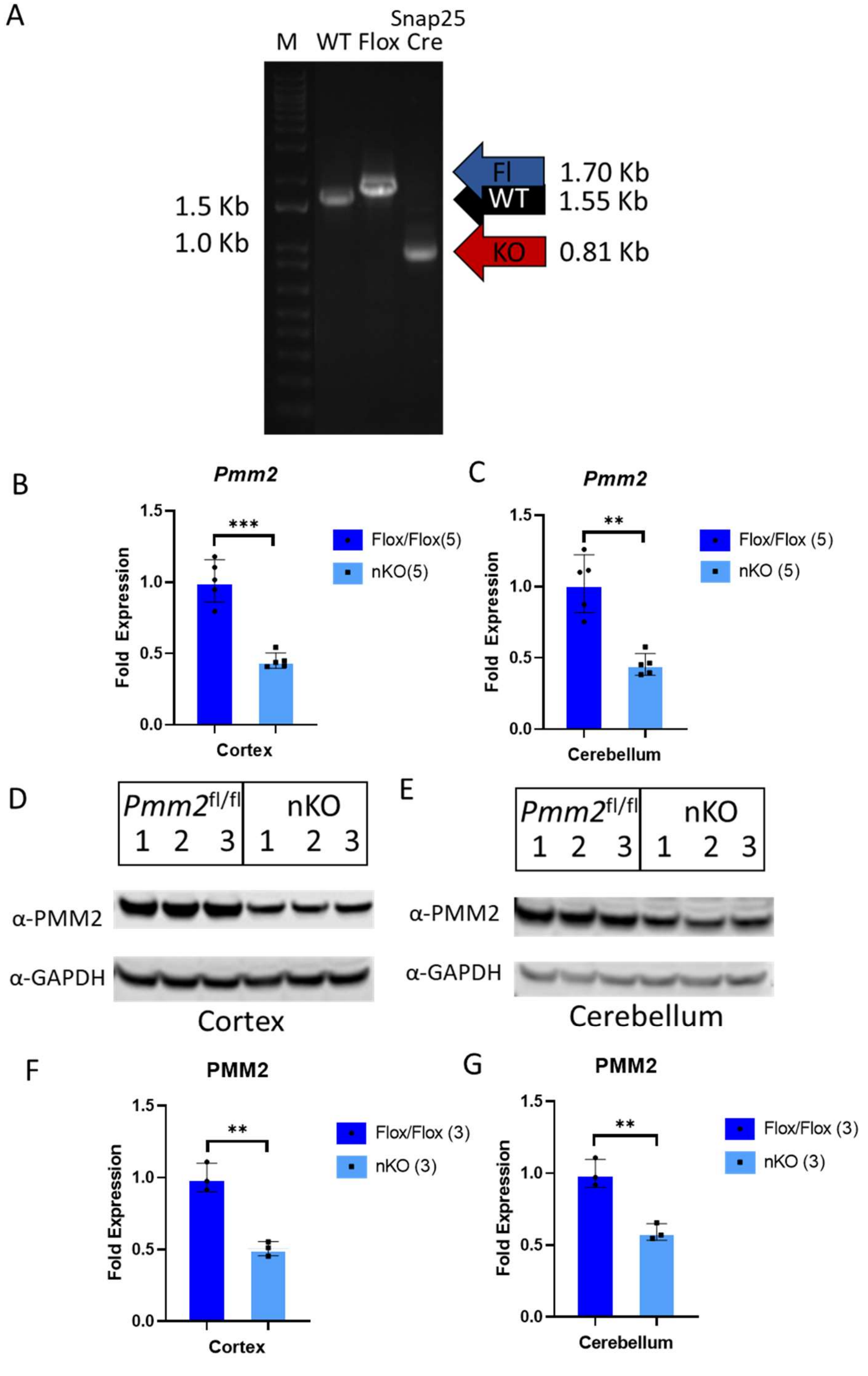


**Supplemental Figure 1. Molecular validation of *Pmm2* nKO mice.** Genomic Recombination Assay PCR results of cortical DNA from WT, Conditional KO (Flox) and nKO (Snap25-Cre) mice. B and C, qPCR quantitation of *Pmm2* expression in cortex and cerebellum, respectfully. D and E, Western for PMM2 protein in cortex and cerebellum, respectfully. F and G, Quantification of Western using GAPDH as a loading control. Data represented as mean ± SD. Welch's t-test. **, P < 0.01; ***, P < 0.001

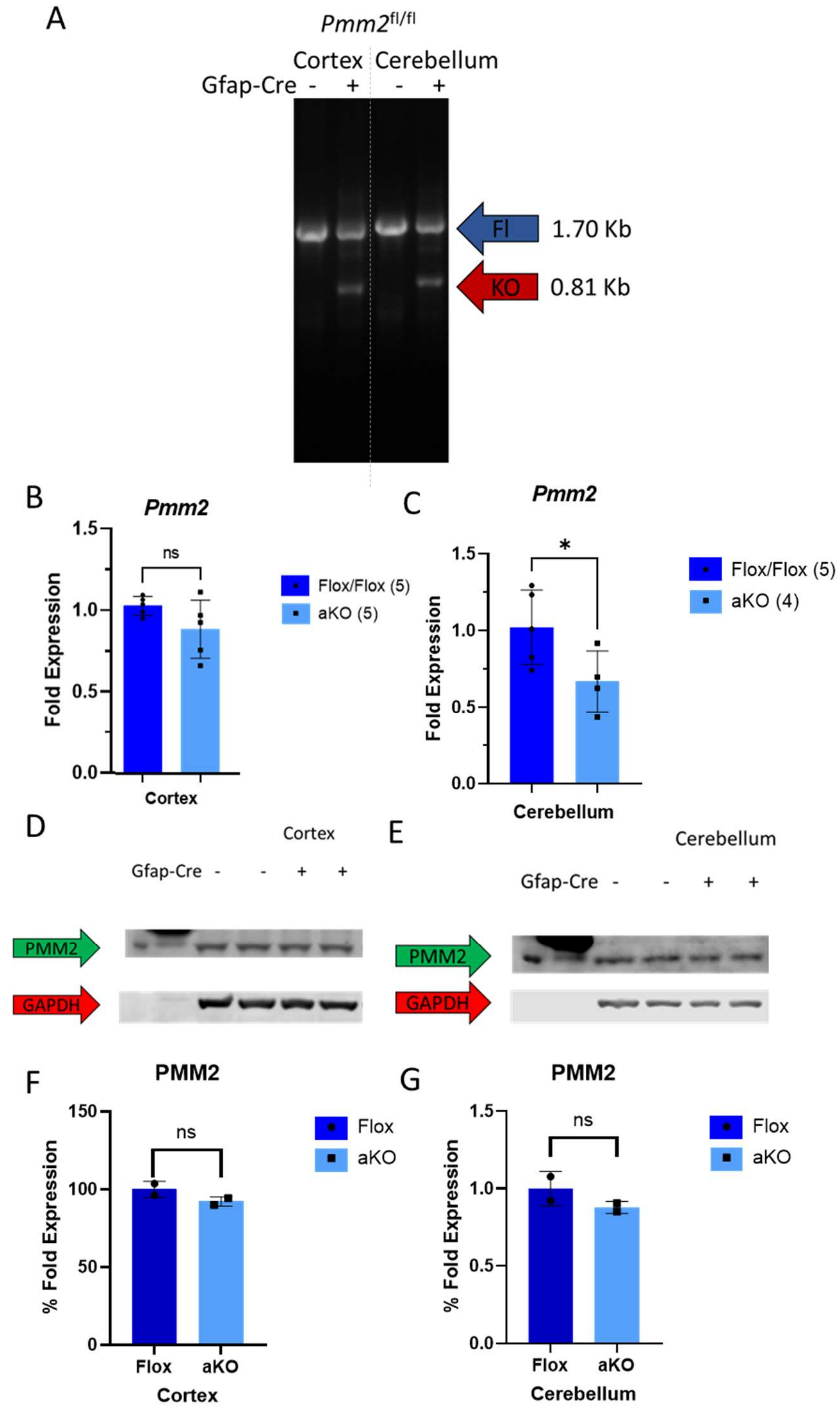


**Supplemental Figure 2. Molecular validation of *Pmm2* aKO mice.** Genomic Recombination Assay PCR results of cortical and cerebellar DNA from Conditional KO and aKO (Gfap-Cre) mice. B and C, qPCR quantitation of *Pmm2* expression in cortex and cerebellum, respectfully. D and E, Western for PMM2 protein in cortex and cerebellum, respectfully. F and G, Quantification of Western using GAPDH as a loading control. Data represented as mean ± SD. Welch's t-test. *, $P < 0.05$

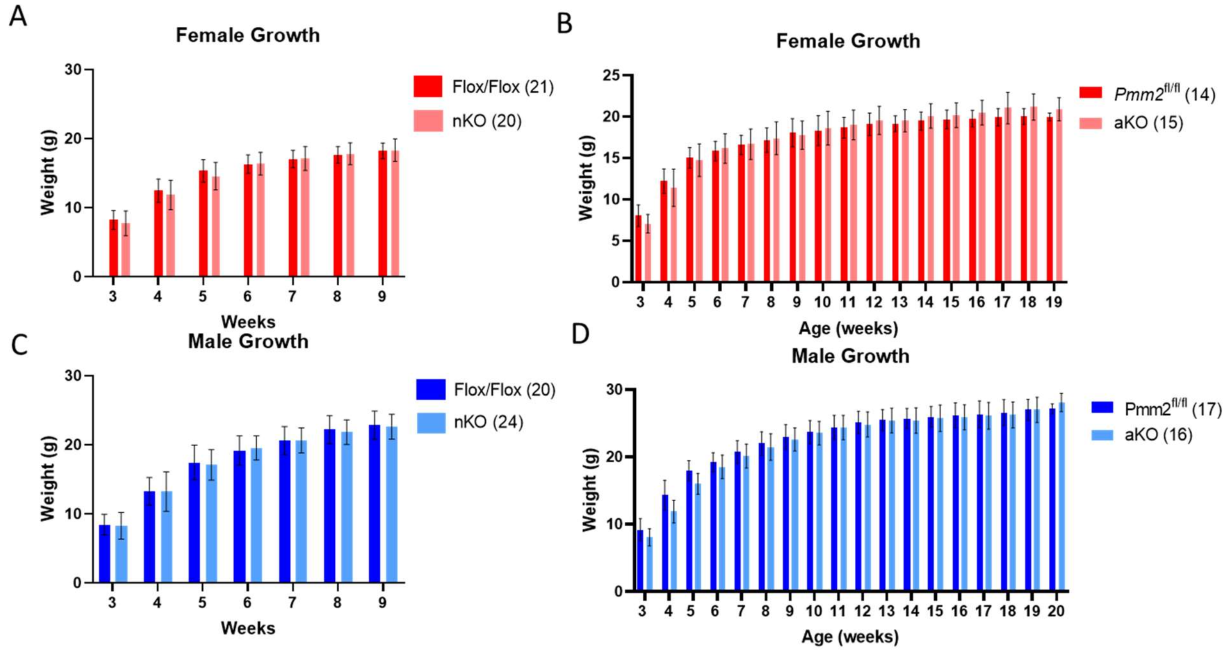


**Supplemental Figure 3. Growth parameters of nKO and aKO mice.** Weekly weights of female (A) nKO and (B) aKO mice and of male (C) nKO and (D) aKO mice. Growth charts initially analyzed using generalized linear mixed-effects model. Pairwise comparisons at each timepoint analyzed using Fisher's Least Significant Differences test. Data represented as mean ± SD.

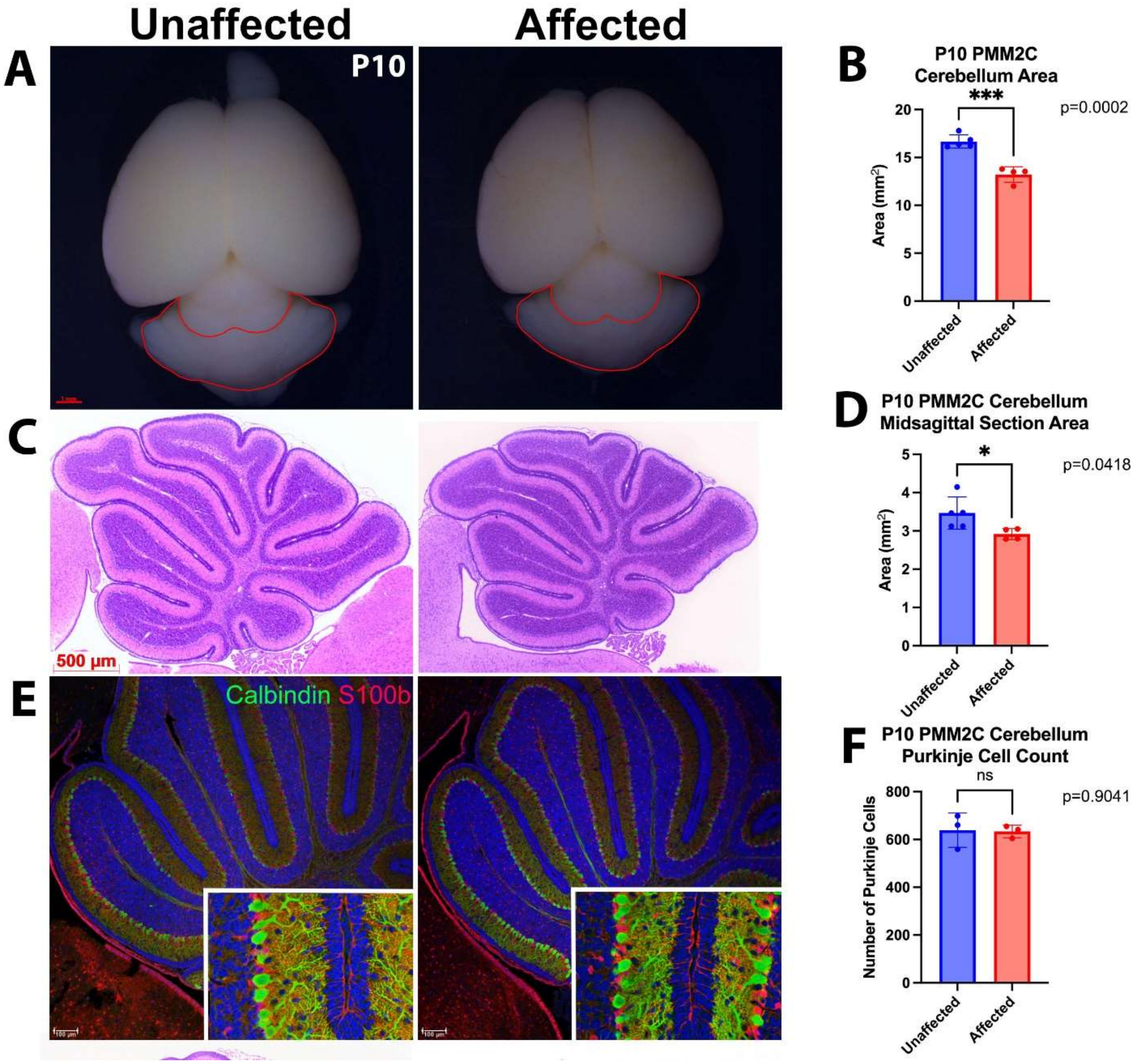


**Supplemental Figure 4. Histologic evaluation of cerebellum in P10 *Pmm2* eKO mice.** A, Cerebellar area of unaffected (*Pmm2*$^{fl/+}$; Nestin-Cre) and affected (*Pmm2*$^{R137H/fl}$; Nestin-Cre) mouse brains at P10. B, Unaffected and affected P10 mice show a significant difference in cerebellum area, *P* = 0.0002, unaffected n=5, affected n=4. C, Representative images of midsagittal sections stained with hematoxylin and eosin of P10 unaffected and affected mice. D, Unaffected and affected P10 mice show a significant reduction in midsagittal area, *P* = 0.0418, unaffected n=5, affected n=4 E, Representative immunohistochemistry images of cerebellum of affected and unaffected mice with staining for Purkinje cells (Calbindin), glia (S100b), and DNA (Hoechst). F, Purkinje cell numbers in the unaffected and affected P10 mice remain unchanged between the two groups. Comparisons analyzed using an unpaired T-test, Data presented as mean ±SD, *,  *P* < 0.05; **, *P* < 0.01; ***, *P* < 0.001

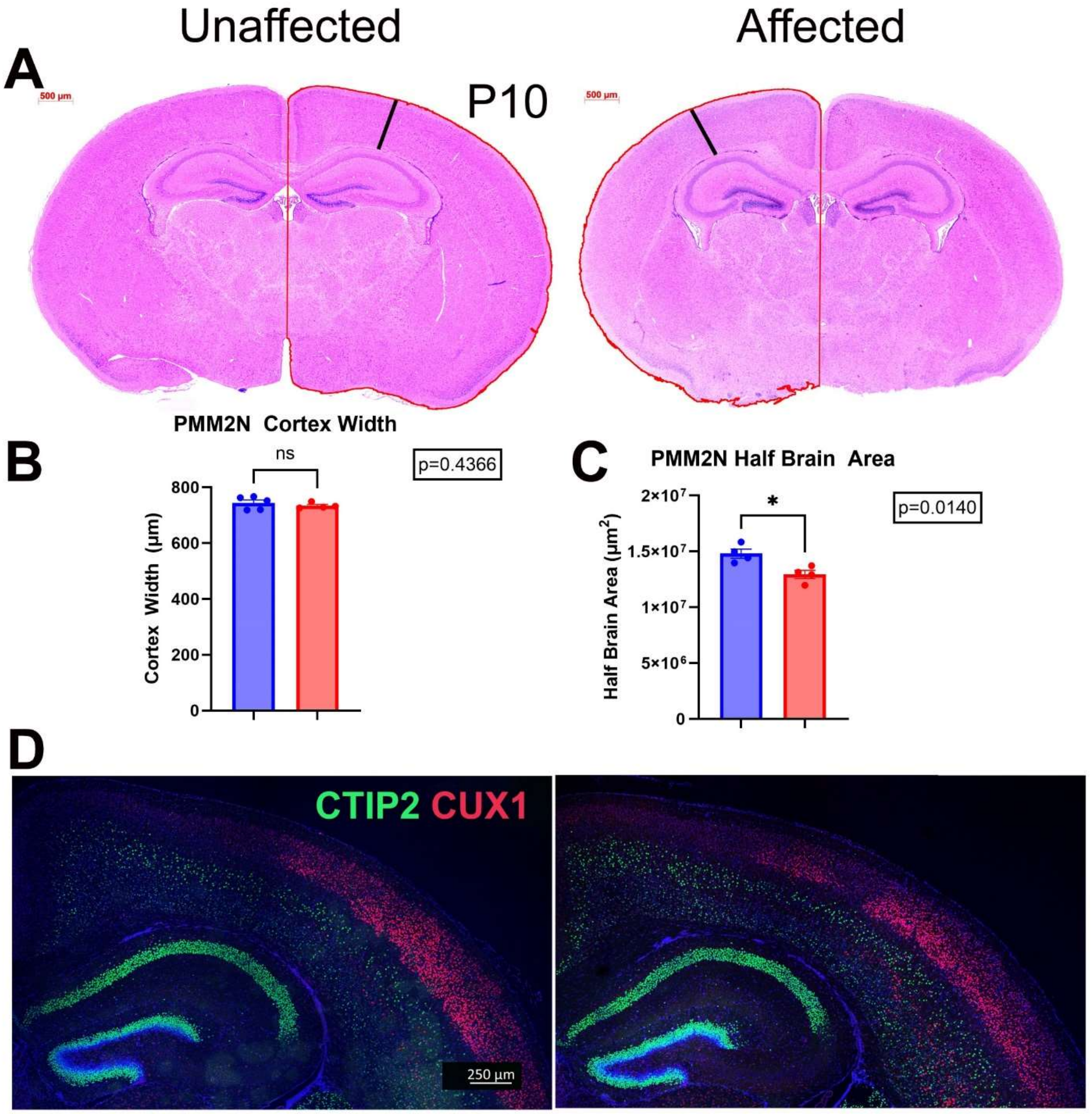


**Supplemental Figure 5. Histologic evaluation of cortex in P10 *Pmm2* eKO mice.** A, Representative images of coronal cortex sections stained with hematoxylin and eosin of P10 unaffected ($Pmm2^{fl/+}$; Nestin-Cre) and affected ($Pmm2^{R137H/fl}$; Nestin-Cre) mice. Red outlines indicate regions used for measurements of hemisphere area; black lines indicate cortical width measurements. B, Unaffected and affected P10 mice displayed no significant difference in cortex width. C, Unaffected and affected P10 mice showed a significant difference in hemisphere area, $P$ = 0.0140, n=4 for each genotype. D, Representative immunohistochemistry images of the cortex of affected and unaffected mice with staining for CTIP2 (layer 5), Cux1 (layer 2-4), and DNA (Hoechst) showed no stark differences. Comparisons analyzed using an unpaired T-test, data represented as mean ±SD,*, $P < 0.05$
